# Neural crest cells bulldoze through the microenvironment using Aquaporin-1 to stabilize filopodia

**DOI:** 10.1101/719666

**Authors:** Rebecca McLennan, Mary C. McKinney, Jessica M. Teddy, Jason A. Morrison, Jennifer C. Kasemeier-Kulesa, Dennis A. Ridenour, Craig A. Manthe, Rasa Giniunaite, Martin Robinson, Ruth E. Baker, Philip K. Maini, Paul M. Kulesa

## Abstract

Neural crest migration requires cells to move through an environment filled with dense extracellular matrix and mesoderm to reach targets throughout the vertebrate embryo. Here, we use high-resolution microscopy, computational modeling, and in vitro and in vivo cell invasion assays to investigate the function of Aquaporin-1 (AQP-1) signaling. We find that migrating lead cranial neural crest cells express AQP-1 mRNA and protein, implicating a biological role for water channel protein function during invasion. Differential AQP-1 levels affect neural crest cell speed, direction, and the length and stability of cell filopodia. Further, AQP-1 enhances matrix metalloprotease (MMP) activity and colocalizes with phosphorylated focal adhesion kinases (pFAK). Co-localization of AQP-1 expression with EphB guidance receptors in the same migrating neural crest cells raises novel implications for the concept of guided bulldozing by lead cells during migration.

## INTRODUCTION

Cell migration is essential during embryogenesis to gastrulate, elongate the vertebrate axis, and distribute cells into the periphery to contribute to organ development. Despite the importance of cell migration to human development and disease, it is still unclear what mechanisms enable cells to invade the dense ECM, mesoderm and other cell types characteristic of the embryonic microenvironment. The complexity of the embryonic microenvironment means that invading cells must rapidly change cell shape and volume, form and sustain protrusions that penetrate different sized gaps, and attach to and remodel the ECM. Thus, there is a tremendous need to identify and test the function of molecules critical to embryonic cell migration and better understand their mechanistic basis.

Neural crest cell migration is one of the most prevalent examples of how cells efficiently distribute throughout the growing vertebrate embryo to precise targets. In the head, cranial neural crest cells must invade through dense ECM, loosely connected mesoderm, and migrating endothelial cells. Yet, it has remained unclear how the migrating neural crest cells that first encounter the embryonic microenvironment penetrate small gaps between mesodermal cells and degrade the ECM to move in a directed manner to peripheral targets. By combining dynamic in vivo imaging and super resolution microscopy with gain- and loss-of-function experiments, we are poised to examine the function of genes presumed critical to neural crest cell migration. Thus, the embryonic neural crest is an attractive in vivo model to study the function of cell invasion genes in mechanistic detail.

Using single cell RT-qPCR and transcriptome profiling, we discovered the enhanced expression of several genes in the most invasive chick cranial neural crest cells, including *AQP-1* (McLennan et al., 2015a; Morrison et al., 2017a). AQP-1 is a transmembrane channel protein that facilitates the flux of water across the plasma membrane (Agre et al., 1993) and is one member of a family of at least 13 aquaporins (Ishibashi et al., 2011). AQP-1 has been detected in several aggressive human cancers and its expression correlates with poor disease prognosis (Tomita et al., 2017; De leso and Yool, 2018). However, the mechanistic basis of AQP-1 function is still unclear since studies have been limited to analyzing cell behaviors using in vitro assays. This has led to the generation of several distinct hypothetical mechanisms of AQP-1 function. For example, AQP-1 is thought to allow cells to rapidly change cell volume to form thin filopodial protrusions that squeeze in between neighboring cells (Papadopoulos et al., 2008; Verkman, 2009; Karlsson et al., 2013) or collapse a cell protrusion to retreat from a repulsive signal (Cowan et al., 2000). Alternatively, AQP-1 may function to stabilize a filopodium since aquaporins have been shown in vitro to localize to the front end of migrating cells (Saadoun et al., 2005) and are speculated to function as a cell motility engine (Condeelis, 1993; Stroka et al., 2014). Thus, a better understanding of the in vivo function of AQP-1 and its connection to cell guidance signaling would be beneficial to studies in cell migration and invasion in cancer and developmental biology.

In this study, we examined the expression and function of AQP-1 during chick cranial neural crest cell migration and possible upstream guidance and downstream AQP-1 effectors. We first characterized *AQP-1* mRNA and AQP-1 protein expression within migrating cranial neural crest cells using 3D confocal and super resolution microscopy. We examined the role of AQP-1 in vitro and in vivo by measuring changes to the neural crest cell migratory pattern, individual cell behaviors, and filopodial dynamics after gain- and loss-of-function of AQP-1. To test our hypothesis of a “bulldozer” role for AQP-1 expressing neural crest cells, we analyzed focal adhesion activity, integrin localization and ECM degradation after AQP-1 manipulation. We examined the possible co-localization of Eph guidance receptor expression with AQP-1 in the same migrating neural crest cells using multiplexed fluorescence in situ hybridization. Lastly, we use computer simulations to predict what AQP-1 related functions are likely to enhance cell migration. Together, our data demonstrate a critical role for AQP-1 during cranial neural crest cell migration and offer a mechanistic basis for in vivo AQP-1 function to promote cell invasion.

## RESULTS

### *Aquaporin-1* mRNA is expressed in migrating cranial neural crest cells and preferentially within the lead subpopulation

To detect the expression of *AQP-1* mRNA in vivo specifically within migrating cranial neural crest cells, we took advantage of our recently optimized integrated protocol combining RNAscope multiplexed fluorescence in situ hybridization, immunohistochemistry (IHC), and tissue clearing with FRUIT (Morrison et al., 2017b). This enabled the detection of *AQP-1* mRNA expression in migrating HNK-1 labeled neural crest cells (Fig. 1A-C). By spot counting the number of *AQP-1* transcripts per neural crest cell, we found that *AQP-1* expression was visible in migrating cranial neural crest cells observed at Hamburger and Hamilton stage (HH) 13 (Hamburger and Hamilton, 1951) enroute from rhombomere 4 (r4) to the second branchial arch (BA2) (Fig. 1C; neural crest cells are color-coded to show number of *AQP-1* counted transcripts). In a typical r4 neural crest cell migratory stream, the highest numbers of detected transcripts of *AQP-1* were found to be in the cell subpopulation at the front of the migratory stream (Fig. 1C). We determined this by quantifying the fluorescence signal of *AQP-1* mRNA expression as a function of distance along the migratory stream (from the neural tube towards BA2) in a large number of cells/embryos (Fig. 1D). This confirmed that lead neural crest cells showed significantly higher level of *AQP-1* expression compared to cells further back in the invading stream (Fig. 1D).

**Figure 1:**
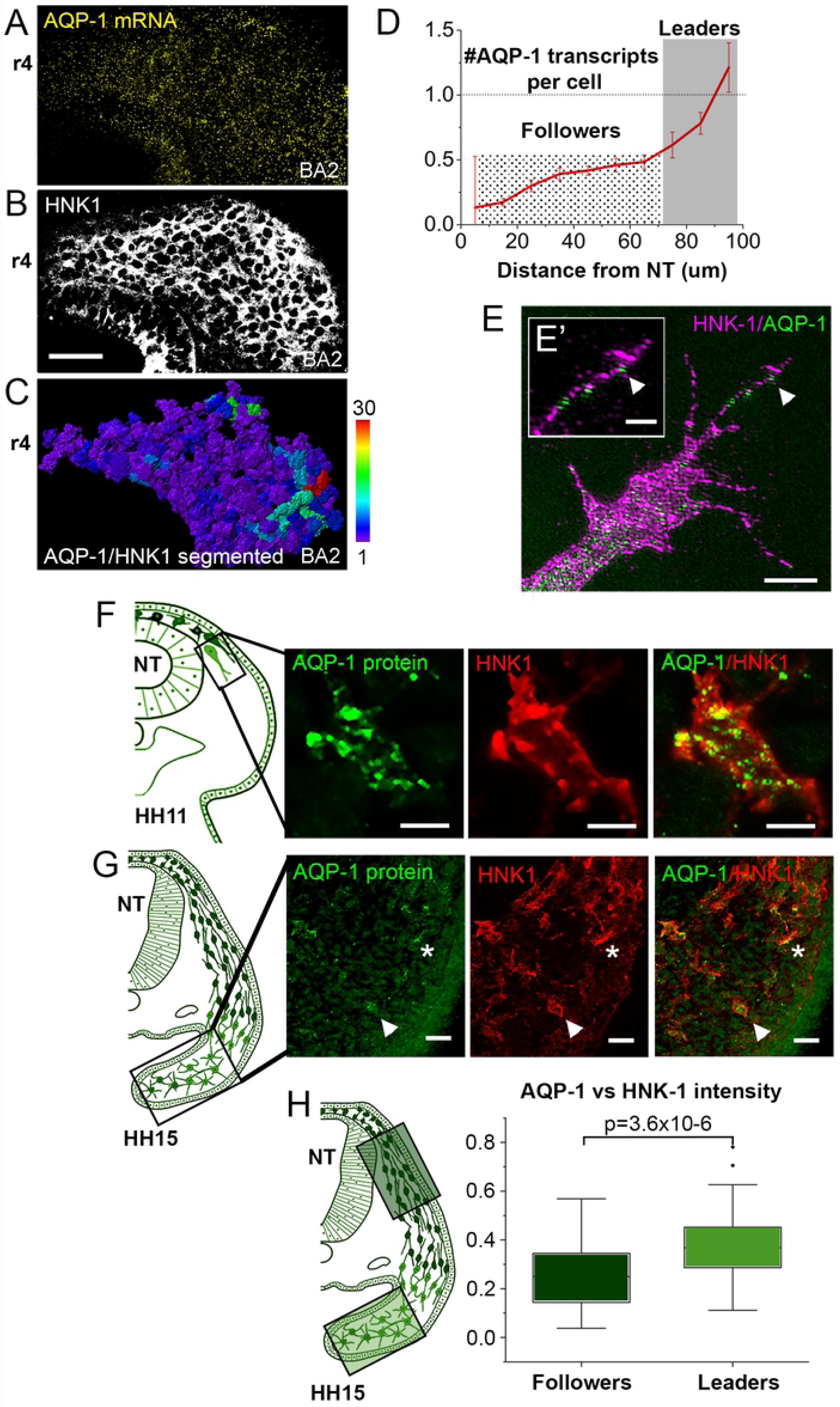
Aquaporin-1 is expressed by neural crest cells at the invasive front of the migratory stream and it located in filopodia. (A-C) Expression of *AQP-1* mRNA by RNAscope (B) in migrating cranial neural crest cells co-labeled with HNK-1 (A) with number of *AQP-1* transcripts per neural crest cell counted (C). Bar=30 um. (D) Quantification of the number of *AQP-1* mRNA transcripts per cell versus distance from the neural tube. n=∼7500 cells, n=10 embryos. (E) High resolution image of the neural crest cell protrusions in vitro showing AQP-1 protein (green) in the filopodia (arrowheads). Bar=5 um and 2 um for inset. (F) Schematic representation of transverse section and high resolution image of a migrating neural crest cell (red) at HH11 with AQP-1 protein (green). Bar=10 um. (G) Schematic representation of transverse section and migrating neural crest cells (red) at HH15 with AQP-1 protein (green). AQP-1 rich neural crest marked with arrowhead and asterisk. Bar=20 um. (H) Quantification of the AQP-1 protein intensity versus HNK-1 intensity from transverse cryostat sections, n=70 leader cells, n=51 follower cells.

### Aquaporin-1 protein is localized on the cell membrane of neural crest cells, including filopodia in vitro and expressed throughout the migratory stream in vivo

Whether AQP-1 protein is expressed within subregions of individual migrating neural crest cells is difficult to determine since it is challenging to resolve fluorescence signal in thin (1-2 um wide) protrusions. Structured illumination microscopy (SIM) has emerged as an excellent tool to resolve diffraction limited issues by increasing optical resolution (Gustafsson et al., 2008). When we applied SIM to visualize AQP-1 protein location within migrating neural crest cells in vitro, we discovered AQP-1 was present on the cell membranes including the tips of filopodia (Fig. 1E). To determine AQP-1 protein expression in vivo, we analyzed AQP-1 protein by IHC within HNK-1 labeled migrating neural crest cells and confirmed AQP-1 expression throughout the migratory stream (Fig. 1F, G; HH11-15). Further, by quantifying fluorescent intensity in individual leader and follower neural crest cells, we determined that there is a higher level of AQP-1 protein in the lead neural crest cells in vivo (Fig. 1H). These data suggested that AQP-1 may be influencing the formation, stability and/or the retraction of neural crest cell filopodia at the leading edge of the migratory stream.

### Aquaporin-1 perturbations in vitro alter neural crest cell speed

To begin to determine the function of AQP-1 in cranial neural crest cell migration, we explanted cranial neural tubes in an in vitro assay and measured changes in cell migratory behaviors with confocal time-lapse microscopy (Fig. 2A, B). We took advantage of Acetazolamide (AZA), a chemical inhibitor of AQP-1 function (Bin and Shi-Peng, 2011; Zhang et al., 2012; Cai et al., 2018; Ameli et al., 2012). Although AZA can also affect AQP-4 function, AQP-1 is the only aquaporin significantly expressed by the invasive front of the migratory stream and therefore the only aquaporin significantly affected by AZA in this assay (Huber et al., 2007; McLennan et al., 2015a; Morrison et al., 2017a). When we inhibited AQP-1 function in vitro, we observed that the migrating neural crest cells moved significantly slower than control neural crest cells (in the presence of DMSO) (Fig. 2B, C; Supplemental Movie 1). Representative tracks show that neural crest cells exposed to AZA remained closer to the neural tube explant and other cells than controls (Fig. 2E).

**Figure 2:**
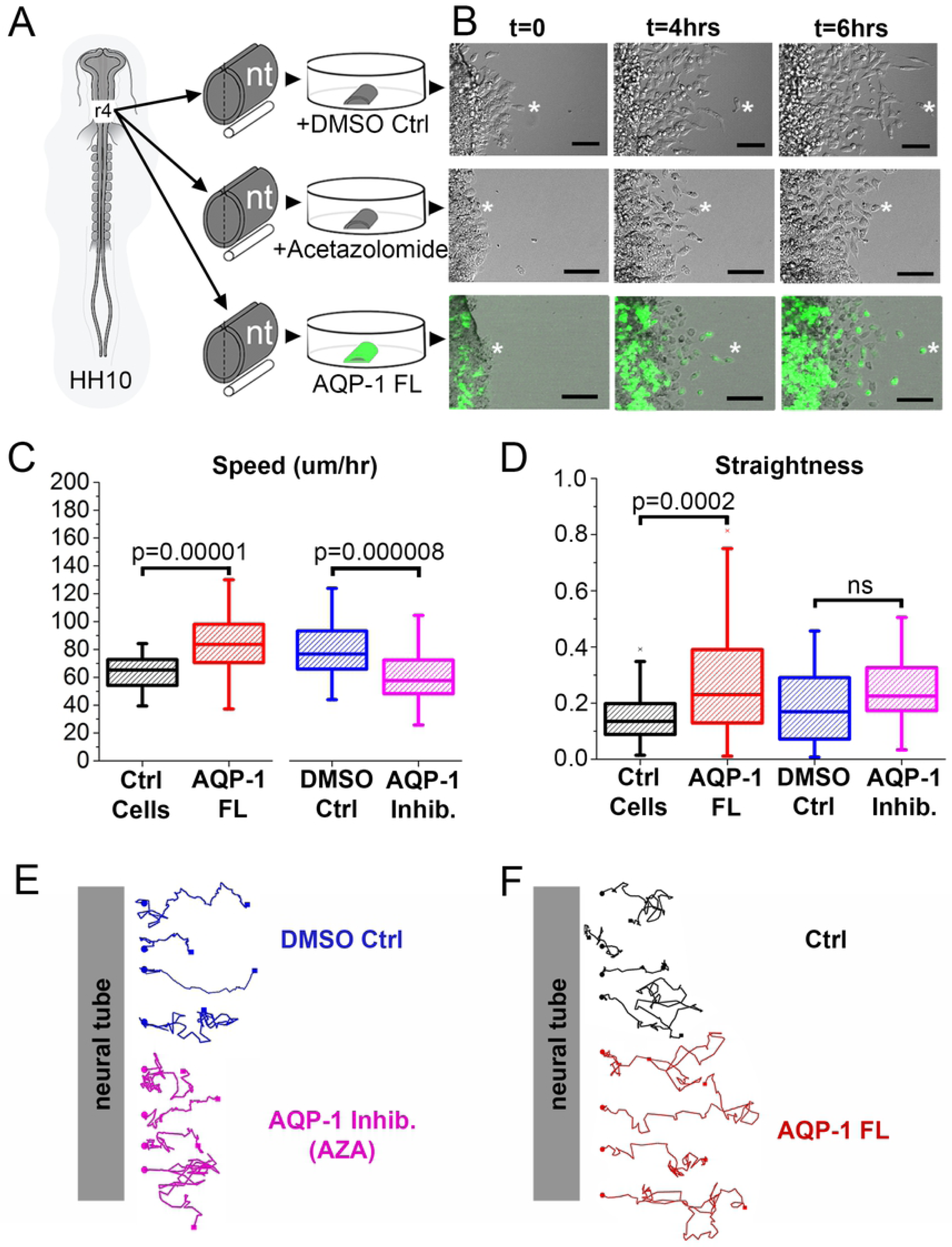
AQP-1 modifies neural crest cell speed in vitro. (A) Schematic representation of experimental setup. (B) Representative images of in vitro neural crest cell time-lapses with neural crest cells exposed to DMSO (control for AZA treatment), AZA and transfected with AQP-1 FL. Bar=50 um. (C) Box plot of the Speed (microns/hr) of neural crest cells with overexpression of AQP-1 (red, n=70 cells) and corresponding controls (black, n=44 cells), AQP-1 inhibition (pink, n=57 cells) and corresponding controls (blue, n=43 cells). (D) Box plot of the Straightness of neural crest cells with overexpression of AQP-1 (red, n=71 cells) and corresponding controls (black, n=31 cells), AQP-1 inhibition (pink, n=26 cells) and corresponding controls (blue, n=27 cells). (E) Representative tracks of neural crest cells with AQP-1 inhibition (pink) and corresponding DMSO controls (blue). Start points are marked with circles, end points are marked with squares. (F) Representative tracks of neural crest cells with overexpression of AQP-1 (red) and corresponding controls (black). Start points are marked with circles, end points are marked with squares. nt, neural tube; AZA, Acetazolamide; ctrl, control; t, time; hrs, hours.

To determine changes in cell migration after AQP-1 gain-of-function, we over-expressed AQP-1 in premigratory neural crest cells (by transfection with AQP-1 FL) and again explanted neural tubes in culture (Fig. 2A, B). AQP-1 over-expression resulted in neural crest cells that moved significantly faster when compared to non-transfected cells in the same culture (Fig. 2B, C; Supplemental Movie 1). Neural crest cells overexpressing AQP-1 were also significantly faster when compared to EGFP only (pMES) transfected cells from different cultures that were prepared and imaged concurrently (Supplemental Fig. 1D). Typical cell tracks from gain-of-function of AQP-1 experiments confirmed neural crest cells moved further away from the neural tube explant in the same amount of time (Fig. 2F). Interestingly, when AQP-1 was overexpressed, cells migrated with increased straightness compared to their controls (Fig. 2D). These in vitro data suggested that AQP-1 significantly influences the speed at which neural crest cells migrate.

### Inhibition of Aquaporin-1 in vivo results in fewer neural crest cells invading branchial arch 2

To determine the in vivo function of AQP-1, we first knocked down AQP-1 function in premigratory cranial neural crest cells by morpholino (MO) transfection. Similar to in vitro experiments, AQP-1 MO transfected neural crest cells did not travel as far as in control MO transfected neural crest cells (Fig. 3A, C, D). In AQP-1 MO transfected embryos, we found fewer migrating transfected neural crest cells in BA2 compared to control MO transfected embryos (Fig. 3B). To further verify the in vivo loss of AQP-1 function in neural crest cell migration, we microinjected AZA into the paraxial mesoderm adjacent to r4, prior to neural crest cell exit from the dorsal neural tube. AZA injected sides of embryos showed fewer migrating neural crest cells migrating into BA2 compared to control sides of the same embryos (Fig. 3F) and control DMSO injected embryos, as seen with immunolabeling of neural crest cells using HNK-1 (Fig. 3G). We also measured fewer neural crest cells present in BA2 when injected with AZA as quantified by percentage of HNK-1 immunostaining signal in BA2 (Fig. 3E). The furthest distance migrated after AZA injection was the same as control, either due to cells escaping AZA, dilution/degradation of the AZA over time or differences in how AZA versus a morpholino affect AQP-1 activity (Supplemental Fig. 1E). There was no significant difference of HNK-1 immunostaining in the control embryos when comparing control side to DMSO injected side (Supplemental Fig. 1F). These data suggested that AQP-1 expression, and therefore function, promotes efficient neural crest cell invasion in vivo.

**Figure 3:**
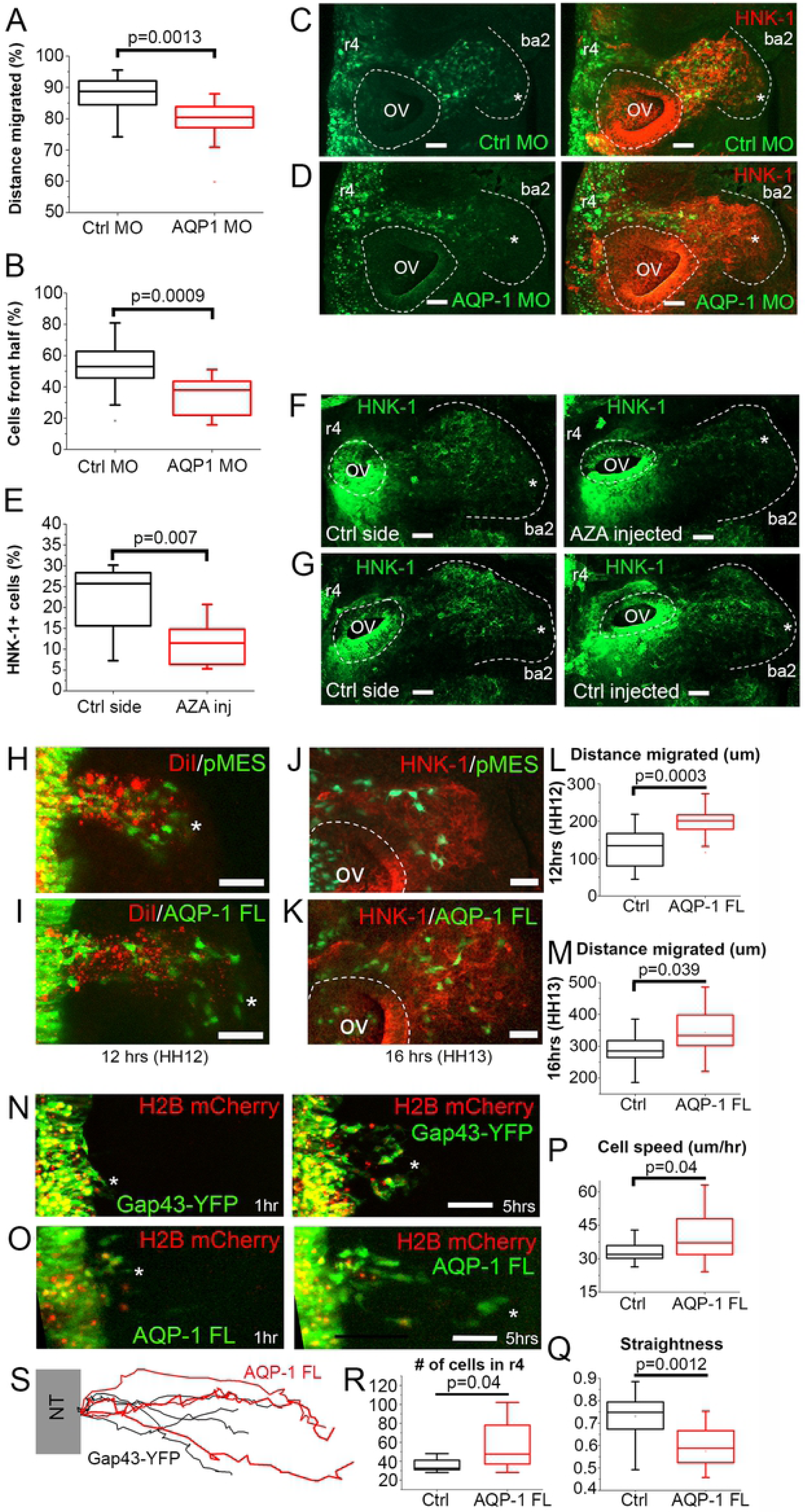
Neural crest cell migration is decreased when AQP-1 is knocked down and increased when AQP-1 is overexpressed in vivo. (A) Box plot of the distance migrated by neural crest cells as a percentage of the whole stream, n=6 embryos per treatment. (B) Box plot of the percent area of the branchial arch (front 50% of the stream) containing HNK-1 positive neural crest cells, n=6 embryos per treatment. (C) HH14 embryo in which neural crest cells (red) are transfected with control MO (green). Bar=50 um. (D) HH14 embryo in which neural crest cells (red) are transfected with AQP-1 MO (green). Bar=50 um. (E) Box plot of the percentage of neural crest cells that migrate into the branchial arches, n=15 embryos per treatment. (F) HNK-1 label of HH15 embryo after injection of AZA into the paraxial mesoderm, control and injected sides shown. Bar=50 um. (G) HNK-1 label of HH15 embryo after control injection of DMSO into the paraxial mesoderm, control and injected sides shown. Bar=50 um. (H) Neural crest migration 12 hours after premigratory neural crest cells were labeled with DiI (red) and transfected with pMES (green). *, end of stream. Bar=50 um. (I) Neural crest migration 12 hours after premigratory neural crest cells were labeled with DiI (red) and transfected with AQP-1 FL (green). *, end of stream. Bar=50 um. (J) Neural crest migration 16 hours after premigratory neural transfected with pMES (green) and then labeled with HNK-1 (red). Bar=50 um. (K) Neural crest migration 16 hours after premigratory neural transfected with AQP-1 FL (green) and then labeled with HNK-1 (red). Bar=50 um. (L) Box plot of the distance migrated by control (pMES) and AQP-1 FL transfected neural crest cells 12 hours after transfection, n=16 pMES embryos, n=15 AQP-1 FL embryos. (M) Box plot of the distance migrated by control (pMES) and AQP-1 FL transfected neural crest cells 16 hours after transfection, n=11 pMES embryos, n=15 AQP-1 FL embryos. (N) Selected images from a time-lapse showing neural crest cells migrating into paraxial mesoderm transfected with a nuclear marker (H2B mCherry, red) and Gap43-YFP (green). *, most invasive cell. Bar=20 um. (O) Selected images from a time-lapse showing neural crest cells migrating into paraxial mesoderm transfected with a nuclear marker (H2B mCherry, red) and AQP-1 FL (green). *, most invasive cell. Bar=20 um. (P) Box plot of the speed of neural crest cells migrating at the front of the stream into paraxial mesoderm, n=16 cells for control, n=11 cells overexpressing AQP-1. (Q) Box plot of the directionality of the same cells tracked in (P). (R) Box plot of the number of transfected cells (pMES and AQP-1 FL) in the neural crest stream 16 hours after transfection, n=8 pMES embryos, n=10 AQP-1 FL embryos. (S) Representative tracks of neural crest cells transfected with Gap43-YFP (black) and AQP-1 FL (red). FL, full length; OV, otic vesicle; ba, branchial arch; AZA, Acetazolamide; ctrl, control; MO, morpholino.

### Overexpression of Aquaporin-1 in vivo enhances neural crest cell invasion

To further investigate the role of AQP-1 in neural crest migration in vivo, we overexpressed AQP-1 (via transfection with AQP-1 full length (FL) construct) in premigratory cranial neural crest cells. Overexpression of AQP-1 caused neural crest cells to migrate further than control pMES-EGFP transfected neural crest cells at both 12 hours (Fig. 3H, I) and 16 hours (Fig. 3J, K) after transfection. Measurements confirmed a significant increase in the distance by AQP-1 FL transfected neural crest cells migrated when compared to pMES transfected neural crest cells analyzed at the same time points (Fig. 3L, M). To determine whether the increased distance migrated when AQP-1 was overexpressed was due to an increase in cell speed, we performed in vivo time-lapse confocal imaging of the migratory streams with five-minute intervals after control (Gap43-YFP) and AQP-1 FL transfections (Fig. 3N, O; Supplemental Movie 2). Cell tracking confirmed that neural crest cells migrated further and faster within the invasive front (front 20% of the migratory stream) when transfected with AQP-1 FL as compared to controls (Fig. 3P, S; Supplemental Movie 2). Interestingly, when neural crest cells were transfected with AQP-1 FL, they also displayed a decrease in cell directionality when compared to pMES transfected neural crest cells (Fig. 3Q). To indicate whether AQP-1 may be playing a role in delamination, the number of transfected cells was counted at 16 hours. Although there were statistically more AQP-1 transfected neural crest than pMES, the range of cell numbers varied greatly and may be due to differences in transfection efficiency as opposed to a role in delamination (Fig. 3R). Together, these data suggested that AQP-1 is critical to the directed neural crest cell invasion but they do not reveal the mechanistic basis underlying AQP-1 function was unclear.

### Aquaporin-1 stabilizes neural crest cell filopodia

To begin to address the mechanistic basis of AQP-1 function during cranial neural crest cell migration, we first perturbed AQP-1 function and used spinning disk confocal microscopy on the lead neural crest cells in vivo to observe any rapid changes in cell morphology and behavior. Specifically, we collected z-stacks at 20 to 30 second intervals of neural crest cells transfected with Gap43-mTurquoise2 as well as AQP-1 MO (knockdown), AQP-1 FL (overexpression) or pMES (control) (Fig. 4A, B, J; Supplemental Movie 3). Images of transfected cells were first processed by creating a binary mask of single cells in ImageJ using the membrane bound Turquoise2 signal and then using the CellGeo software (Tsygankov et al., 2014) and additional calculations to identify and measure filopodia length, position, angle and survival time (Fig. 4C-E). With knockdown of AQP-1, we find a significant reduction in the number and length of filopodia in migrating neural crest cells (Fig. 4F, G). In addition, filopodia in cells lacking AQP-1 retracted at significantly faster rates than control cells leading to less stable filopodia with a shorter survival time (Fig. 4H, I). In contrast, lead neural crest cells with an overexpression of AQP-1 resulted in enhanced stable filopodia with reduced protrusion and retraction rates and a longer survival time (Fig. 4H, I). The average number of filopodia per cell was reduced when AQP-1 was overexpressed but their length was unchanged (Fig. 4F, G).

**Figure 4:**
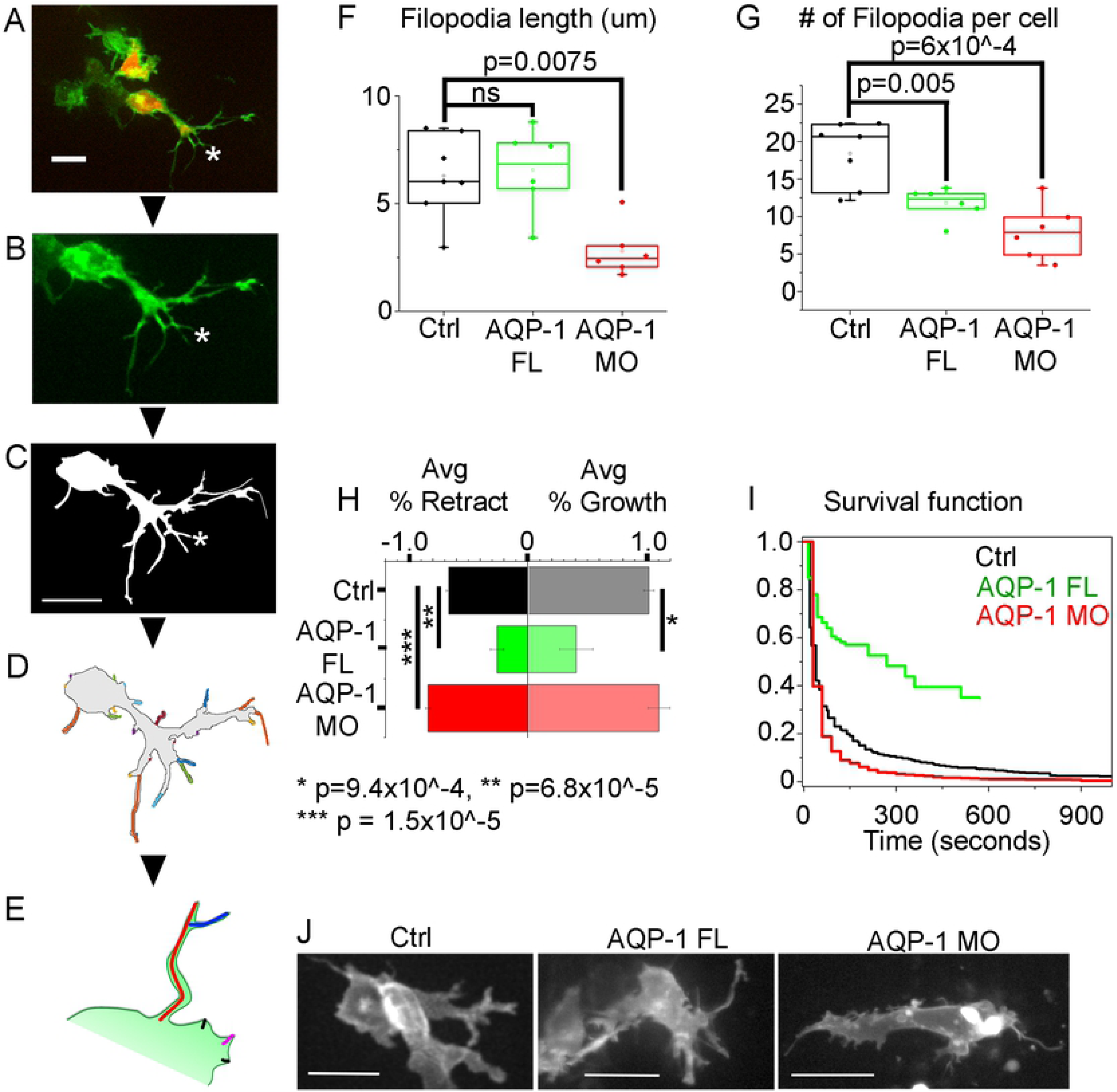
Identification and tracking of in vivo neural crest filopodia reveals AQP-1 expression changes the length, number, growth rate and survival. (A) Front of r4 stream of neural crest cells of a pMES control (red) and Gap43-mTurquoise2 (green) labeled HH13 embryo. (B) mTurquoise2 label of individual cell and (C) masked image of cell for filopodia identification. (D) Each identified filopodium of a single time point for a cell marked in different colors. (E) Schematic of filopodium identification if filopodia are bifurcated; one tip is the main filopodia and the branch constitutes a second filopodium of shorter length. (F) Box plots showing no significant change in filopodia length when AQP-1 is overexpressed but shorter filopodia when AQP-1 is knocked down by morpholino. (G) Box plot of the number of filopodia per cell showing a reduction in number if AQP-1 expression is altered. (H) Percent change in filopodial length from time point to time point for Control, AQP-1 FL and AQP-1 MO cells. AQP-1 overexpression reduces the rate of filopodial growth or retraction but knock down increases only the retraction rate. (I) Survival function for filopodia under three conditions. The probability of a filopodium in a cell overexpressing AQP-1 is much higher than Control. AQP-1 knock down reduces the survival probability of a filopodium. (J) Representative mTurquoise2 membrane labeled cells in vivo. All Bars = 20 um.

We also examined the direction in which the filopodia extended relative to the direction of migration of the cell. A vector was created from the base of the filopodia to the tip and the angle to the direction of motion was calculated (Supplemental Fig. 2C). The control filopodia were roughly bi-polar in distribution, pointing both towards and away from the direction of motion as shown on the rose plot though the Rayleigh test does indicate a uniform distribution (p=0.2) (Supplemental Fig. 2A). With overexpression of AQP-1, the filopodia are found to point randomly in all directions (p=0.22) (Supplemental Fig. 2A). Filopodia after knockdown of AQP-1 by MO on average are directed −60° degrees rostral from the direction of motion roughly towards BA3 (p=4.8×10^-6^) (Supplemental Fig. 2A). We considered the possibility that the longest filopodia may be more important for directional migration by comparing the length of the filopodia found in a 30° window around the direction of motion to the length of filopodia in all other directions. For control cells, the filopodia in the direction of migration are significantly longer than the remaining filopodia (p=5.9×10^-9) (Supplemental Fig. 2B). However, this association is lost in both overexpression and knockdown of AQP-1 and the filopodia in the direction of motion are no longer than filopodia elsewhere in the cell (p=0.31 and p=0.2 respectively) (Supplemental Fig. 2B). These data strongly suggest that AQP-1 functions to stabilize filopodia in migrating neural crest cells; results in the literature suggest that this is likely due to water flux (Papadopoulos et al., 2008; Karlsson et al., 2013; Saadoun et al., 2005).

### Aquaporin-1 influences neural crest cell focal adhesions and ECM degradation

To further explore the role of AQP-1 in neural crest cell invasion, we examined whether AQP-1 is involved in two important aspects of cell migration: cell–ECM attachment and ECM degradation. First, we overexpressed AQP-1 in migrating neural crest cells, isolated those cells from developing embryos and performed RNAseq. The sequencing results were compared to neural crest cells transfected with pMES only. Significant pathways affected by AQP-1 expression include many guidance signaling pathways, such as semaphorins, FGFs, neuregulin and ephrins, as well as the actin cytoskeleton (Fig. 5A). Members of the AQP family have been shown to promote migration by facilitating the recycling of integrins and therefore turnover of focal adhesions that are necessary for cell invasion (Chen et al., 2012). When AQP-1 is overexpressed, *integrins A8* and *B5* were significantly increased (Supplemental Table 1). Therefore, we performed IHC for HNK-1, AQP-1 and phosphorylated FAK on cranial neural crest cells in migrating r4 neural crest in vivo. Confocal imaging with Airyscan detection reveals a colocalization between the puncta of pFAK and AQP-1 IHC (Pearson’s coefficient 0.74), indicating a very close physical proximity of these two proteins (Fig. 5C). To further investigate the possible link of AQP-1 with pFAK and integrins, we performed IHC for HNK-1, pFAK and integrin B1 after transfection with AQP-1 FL (overexpression) or pMES (control) in vitro. Using Airyscan confocal microscropy and spot detection analysis, AQP-1 overexpression leads to less pFAK puncta as well as less integrin B1 puncta on or very near neural crest cell membranes. RNAseq analysis showed that integrin B1 RNA levels did not change when AQP-1 was overexpressed (Supplemental Table 1). Therefore, this result suggests increased turnover of integrin B1 via internalization, which would be consistent with previous literature (Chen et al., 2012), or dissociation of the integrin heterodimers as well as internalization or loss of phosphorylation of FAK (Fig. 5D).

**Figure 5:**
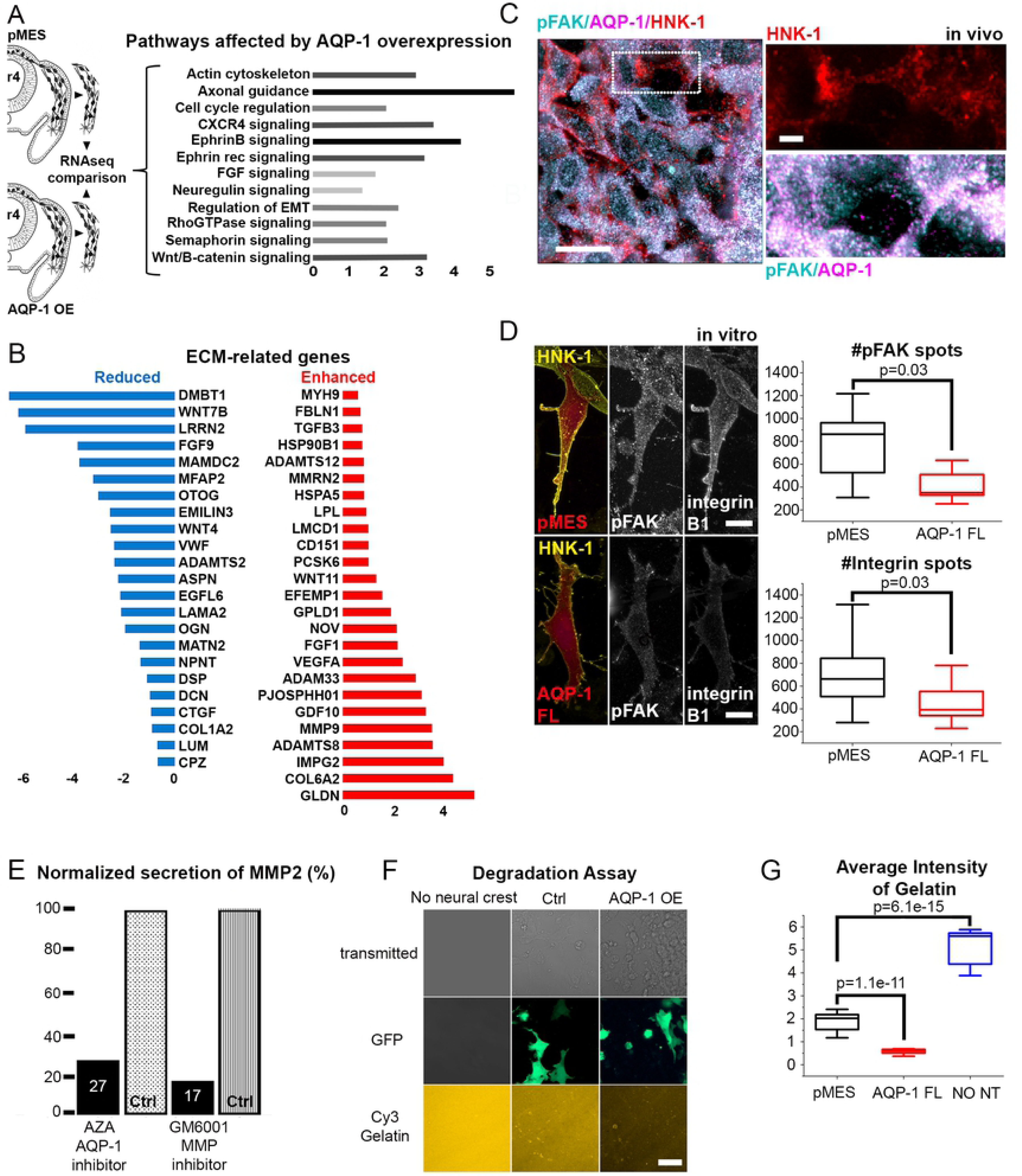
AQP-1 influences focal adhesions and ECM degradation. (A) Integrated pathway analysis displaying developmentally relevant signaling pathways affected by overexpression of AQP-1. (B) Genes associated with ECM GO term that were significantly up- (red) or down-regulated (blue) in response to overexpression of AQP-1. (C) Colocalization of pFAK (cyan), AQP-1 (magenta) and EphB3 (green) protein expression in migrating neural crest cells (HNK-1, red) in vivo. Bar=2 um. (D) High resolution image of neural crest cells (HNK-1, yellow) in vitro transfected with pMES or AQP-1 FL (red) showing pFAK and integrinB1 expression (white). Bar=5 um. Box plot of the number of pFAK spots detected in neural crest cells transfected with pMES or AQP-1 FL, n=10 and 12 respectively. Box plot of the number of integrinB1 spots detected in neural crest cells transfected with pMES or AQP-1 FL, n=10 and 11 respectively. (E) Quantification of in gel zymography using AQP-1 and MMP inhibitors, showing 27% and 17% of MMP activity respectively, n=39 neural tubes each for AZA and control, n=32 neural tubes each for GM6001 and control. (F) In vitro degradation assay. Chambered slides were coated with Cy3-labeled gelatin (orange). Neural tubes transfected with pMES or AQP-1 FL (GFP) were plated onto the gelatin and incubated for 40 hours. No neural tubes were used as baseline control. (G) Box plot of the average intensity of Cy3-labeled gelatin after 40 hours of incubation with no neural crest, pMES transfected neural crest or AQP-1 FL transfected neural crest, n=15 40x images analyzed per condition from n=3 replicates each.

From the RNAseq data, we also focused our attention on ECM related genes as AQP-1, AQP-3 and AQP-4 have all been shown to be positive regulators of the ECM degrading molecules, MMPs (Chen et al., 2015; Ding et al., 2011; Jiang et al., 2014; Wei and Dong, 2015; Xiong et al., 2017; Xu et al., 2011). When AQP-1 is overexpressed in migrating neural crest cells, *MMP9* as well as members of the ADAMs, including *ADAM33* are significantly increased (Fig. 5B). To test whether AQP-1 is upstream of MMP activity, we measured MMP activity from neural crest cells using an in-gel zymography assay. Specifically, neural tube cultures were exposed to either AZA (AQP-1 inhibitor), GM6001 (MMP inhibitor) or used as controls. The media from these cultures was then extracted to determine MMP activity. Neural crest cells exposed to AZA exhibited a reduction of MMP activity of 73% at a size that most likely matches MMP2 (Fig. 5E; Supplemental Fig. 1G). In comparison, neural crest cells exposed to GM6001 exhibited a reduction of MMP activity of 83%, which was our positive control to show that the assay was functional (Fig. 5E; Supplemental Fig. 1G). This result was verified with an in vitro degradation assay, where neural tubes (transfected with AQP-1 FL or pMES) were plated on top of fluorescently labeled gelatin (Garmon et al., 2018). After 40 hours of incubation, the assays were imaged, and average fluorescent intensity of the gelatin was quantified as a read out of degradation (Fig. 5F). Neural tubes transfected with pMES significantly degraded gelatin compared to control wells where no neural tubes were plated as expected (Fig. 5F, G). Transfection with AQP-1 FL resulted in even higher gelatin degradation, which is consistent with the in gel zymography (Fig. 5F, G). These data suggest that AQP-1 is involved in the promotion of integrin turnover and ECM degradation.

### Aquaporin-1 is colocalizes with EphB1 and EphB3 expression in the same migrating cranial neural crest cells

Previous work in commissural axon guidance in mouse has shown evidence for the co-immunoprecipitation of AQP-1 and Eph receptors, namely EphB2, to form a stable complex within a cell (Cowan et al., 2000). Since Ephs and ephrins are implicated in axon pathfinding, Cowan and colleagues speculated that aquaporins may function in the growth cone to help integrate the guidance information elicited by Eph-ephrin clustering (Cowan et al., 2000). Based on this study, we asked whether the Eph receptors EphB1 and EphB3, previously shown to be highly expressed in the most invasive cranial neural crest cells (McLennan et al., 2015a), were co-localized with AQP-1 expression in the same migrating cranial neural crest cells. To address this, we used multiplexed fluorescence in situ hybridization (RNAscope) to label *AQP-1*, *EphB1*, and *EphB3* mRNA expression and HNK-1 IHC to quantify co-expression of these three mRNAs in cranial neural crest cells in vivo (Fig. 6A, B). As before, the HNK-1 channel was used to segment r4 neural crest from whole mount HH13 embryos (white outlines, Fig. 6A, B) and spots of *AQP-1*, *EphB1* and *EphB3* mRNA transcripts were counted per cell. When we quantified the mRNA expression, we found that *AQP-1* mRNA was only found in cells also expressing *EphB1* and/or *EphB3* mRNA though there were many *EphB1* and/or *EphB3* mRNA positive cells without *AQP-1* mRNA (Fig. 6C). When we quantified the average number of detected transcripts in migrating neural crest cells, we found that cells expressing *AQP-1* mRNA had higher number of *EphB1* and *EphB3* mRNA transcripts than their *AQP-1* negative counterparts (Fig. 6D). By IHC, we can also verify that AQP-1 protein and EphB1/B3 protein co-label the same neural crest cells in vivo (Fig. 6E-H).

**Figure 6:**
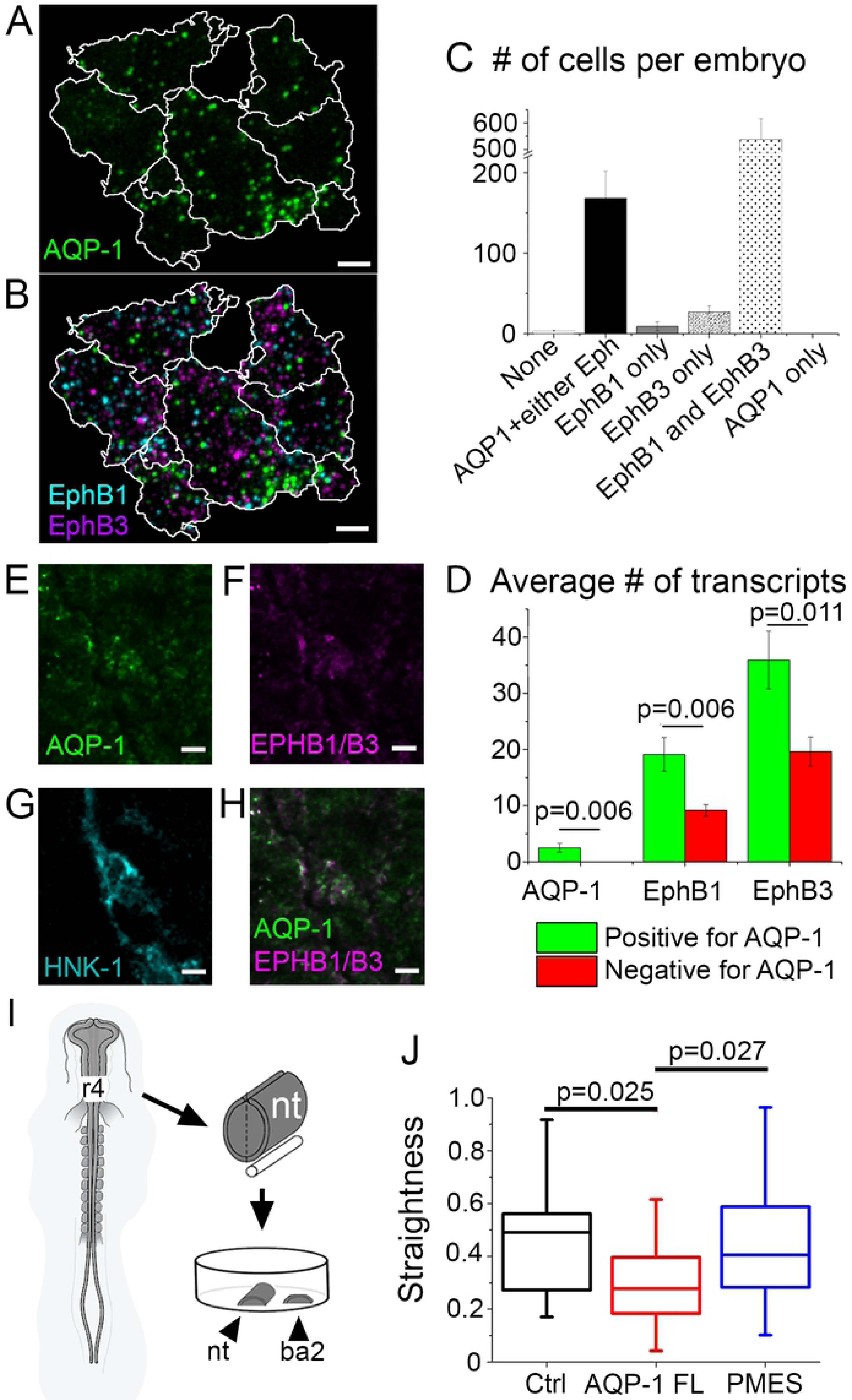
AQP-1 interacts with EphB receptors and is involved in neural crest cell directionality. (A) RNAscope of lead r4 neural crest cells in vivo, HNK-1 signal outlined in white and showing transcripts for *AQP-1* (green). Bar=5 um. (B) RNAscope of same cells in (a) with *AQP-1* (green), *EphB1* (cyan) and *EphB3* (purple). (C) Bar graph of number of cells per embryo expressing different mRNA combinations. (D) Bar graph of average number of transcripts of each gene in cells with or without *AQP-1* mRNA. (E-H) Protein expression of AQP-1 (E), EphB1/B3 (F) and HNK-1 (G) in migrating neural crest cells (H) at HH13. Bar=5 um. (I) Schematic representation of in vitro assay quantified in (J). (J) Box plot of the directionality of neural crest cells responding to branchial arch 2 tissue in vitro, n=18 untransfected cells, n=25 AQP-1 FL cells, n=42 pMES cells.

### Aquaporin-1 is involved in the directed migration of neural crest cells

To further explore the hypothesis that AQP-1 is downstream of guidance, we performed in vitro cultures where migrating neural crest cells were exposed to BA tissue, a known source of guidance (Fig. 6I; Supplemental Movie 4). When AQP-1 was overexpressed in neural crest cells, they were less directed than untransfected controls in the same cultures, or in pMES only transfected neural crest cells (Fig. 6I, J; Supplemental Movie 4). This finding is consistent with the loss of directionality observed when AQP-1 was overexpressed in neural crest cells in vivo (Fig. 3Q), and the loss of polarized filopodia in the direction of migration (Supplemental Fig. 2B). One possible explanation for this result is that by overexpressing AQP-1 motility is increased due to more AQP-1 channels, but these channels are no longer focused by complexing with guidance receptors. Overall, these data suggest that AQP-1 is involved in the fast response of cells to guidance factors, possibly by complexing with guidance receptors, including EphB receptors.

### Computational modeling predicts that enhanced collective cell invasion is achieved by a combination of increased cell speed, filopodia stabilization and ECM degradation

To test hypothetical mechanistic scenarios by which AQP-1 functions during neural crest cell invasion, we used a hybrid computational model of cranial neural crest invasion. The model consists of a discrete, off-lattice model for the cells that is coupled to a continuum, reaction-diffusion model of the dynamics of a known cranial neural crest cell chemoattractant (McLennan et al., 2010; VEGF) on a growing domain. The model is based on our previous studies (McLennan et al., 2015a; McLennan et al., 2012; McLennan et al., 2015b; McLennan et al., 2017) and is fully described in Supplemental Data 1. Briefly, we used a two-dimensional approximation of the neural crest cell migratory domain and incorporated finite size effects by considering cells as hard disks that are not allowed to overlap (Supplemental Table 2; Supplemental Fig. 3). We used a fixed time step model (with a time step of 1 minute) during which a cell senses its environment and moves accordingly. We considered two subpopulations of cells, namely leaders and followers; leaders undertake a biased random walk up a cell-induced gradient of chemoattractant (Supplemental Table 2).

In this study, we modified five parameters, or features, of our model to simulate the mechanisms by which AQP-1 influences cell migration: cell speed; filopodia stability; filopodia polarity; filopodia number and ECM degradation (Fig. 7A). Filopodia stabilization is realized by a leader sampling the microenvironment only every three time steps and moving persistently in between, as opposed to sampling and potentially moving in a different direction at each time step. Filopodia polarization means that a leader only stabilizes its filopodium when a cell makes an informed movement towards a higher concentration of VEGF as opposed to a random movement. Filopodia number is the number of random directions in which a leader samples the concentration of chemoattractant per time step. Since our experimental results showed that overexpression of AQP-1 induced an increase in MMP activity that resulted in ECM degradation (Fig. 5E-G) we introduced into the model the creation of tunnels in the ECM (Fig. 7B). This was implemented by recording the histories of leader positions and defining them as tunnels. Finally, the invasion of cells was assumed to occur on a uniformly growing rectangular domain whose growth was based on biological measurements (McLennan et al., 2015a; McLennan et al., 2012).

**Figure 7:**
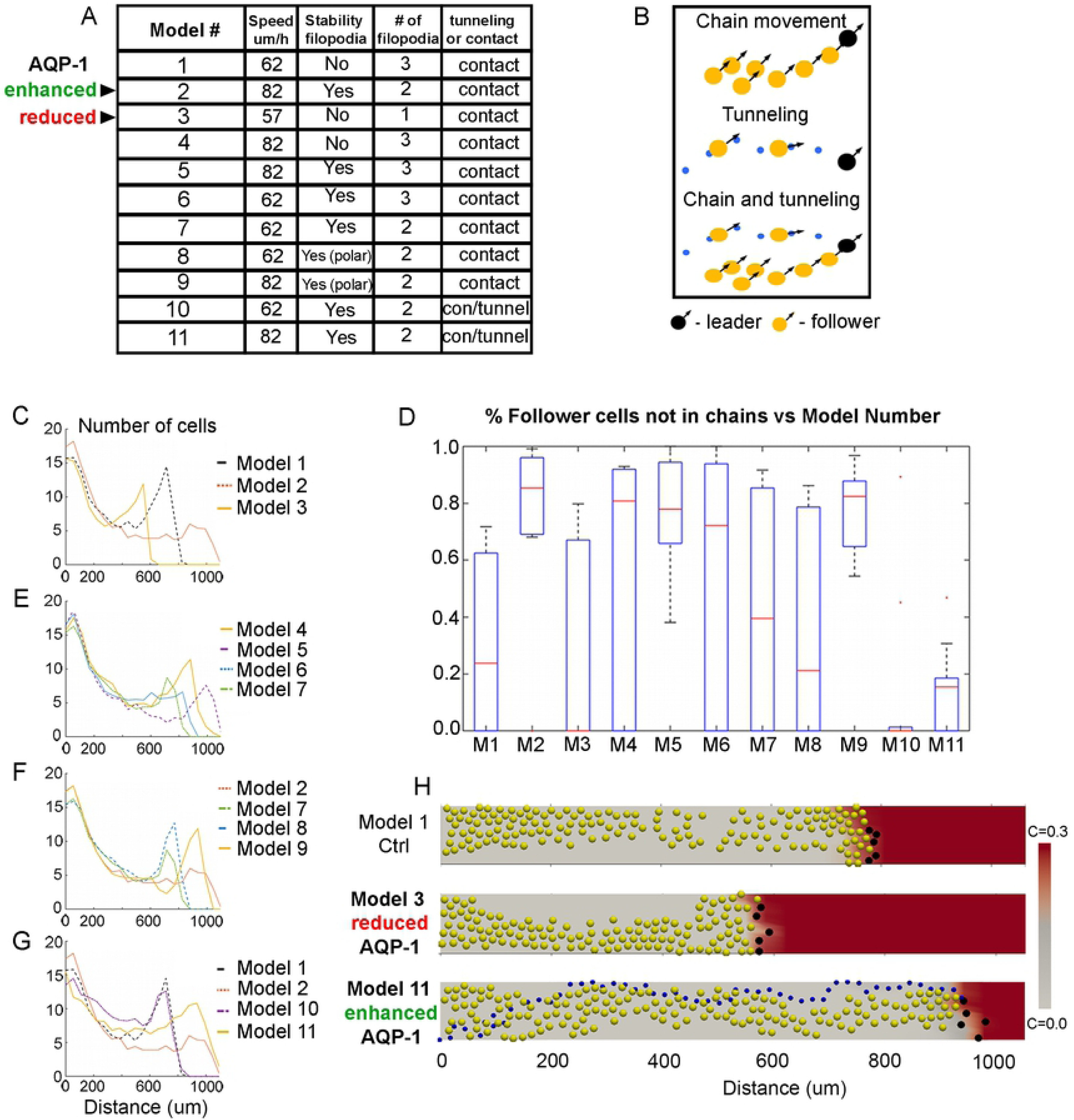
Computer simulations predicts that increased speed, filopodia stabilization and ECM degradation enhance cell invasion. (A) Model parameters for various experimental and hypothetical scenarios. (B) Schematic representations of chain migration where followers (yellow) within a certain distance of each other all adopt the direction of movement of the leader (black), tunnel movement where the tunnel (blue) forged by a leader guides followers that enter the tunnel to move along the tunnel and chain and tunnel movement where a leader can guide followers both via the tunnel it has created and by chain movement. (C, E-G) Distribution of cells along the domain for different models, average of 10 simulations at t=24 hours. (D) Boxplots of fractions of follower cells not in chains (also not in tunnels in Model 10 and Model 11) at t=24 hours. Results are averaged over 10 simulations. For each model, the red line indicates the median, and the bottom and top edges of the box indicate the 25th and 75th percentiles, respectively. The dotted lines extend to the most extreme data points not considered outliers, and the outliers are plotted individually as red dots. M - Model. (H) Snapshots from three selected models at t=24 hours. Black circles – leader cells, yellow circles – follower cells, blue circles – positions of one of the leaders that form a tunnel, c – concentration of chemoattractant (VEGF).

In our computational model we initially quantified changes in the dynamics of cells when we recapitulated the effects of overexpression and downregulation of AQP-1 by modulating relevant model parameters. Different combinations of parameter perturbations are labeled as Model x (x from 1 to 11); the values of parameters for different models can be found in Figure 7A. In model simulations, we measured the density of cells along the two-dimensional migratory domain and the likelihood of migratory stream breakage, defined as the proportion of cells not in chains or tunnels at the end of a simulation (Fig. 7D). In the control case, the cell speed was set to 62 um/hr, filopodia were not stabilized, and there was a representative number of three filopodia per cell (Model 1). To simulate AQP-1 overexpression, the cell speed was increased by approximately 30% to 82 um/hr, filopodia were stabilized and there were two filopodia per cell (Model 2). To simulate inhibition of AQP-1, the cell speed was decreased to 57 um/hr, cells had only a single filopodium, which was not stabilized (Model 3). Initially (Models 1-9), we did not incorporate ECM degradation (i.e. we did not include the tunneling mechanism. For each model, simulations were run ten times with each parameter set, and the results at t=24 hours were averaged to account for random variations. As expected, cells at the front traveled farther when their speed was increased (Fig. 7C; Model 2). Cells traveled shorter distances when their speed was reduced to simulate AQP-1 knockdown (Fig. 7C; Model 3). We did, however, observe an unexpected phenomenon; when the cell speed was increased the simulated cell migratory streams broke apart more frequently and the model did not sustain collective cell migration (Fig. 7D). Migratory stream breakage is not observed in vivo and therefore this observation led us to further investigate cell speed versus filopodia dynamics and the effects upon collective cell migration in silico.

In our simulations, when cell speed was increased, with or without filopodia stabilization, cells at the front traveled farther compared to the control case (compare Model 4 and Model 5 with Model 1) but stream breakage was much more likely to occur (Fig. 7D, E). The number of filopodia (two or three) affected the likelihood of stream breakage at control cell speeds, with more filopodia resulting in a higher likelihood of breakage but with little impact upon the distance migrated (compare Model 6 with Model 7) (Fig. 7D, E). At higher speeds, whether cells had two or three filopodia did not significantly affect the probability of stream breakage or the distance the cells migrated (compare Model 2 with Model 5) (Fig. 7C-E). Stabilization of filopodia increased the farthest distance traveled by cells regardless of the cell speed. It also noticeably reduced the likelihood of stream breakage at control cell speeds, with a smaller effect at enhanced speeds (compare Model 1 with Model 6, and Model 4 with Model 5) (Fig. 7D, E). Additionally, polarization of filopodia had an insignificant effect on the migration pattern (compare Model 7 with Model 8 and Model 2 with Model 9) (Fig. 7D, F).

Since our experimental findings showed that increasing AQP-1 activity led to an increase in MMP activity, and thus to an increase in degradation of the ECM, we included the formation of tunnels by leader cells in silico. We found this had no significant effect on distance the cells migrated, however it did dramatically decrease the percentage of follower cells not in chains to zero (compare Model 1 with Model 10) (Fig. 7D, G). In contrast, to simulate overexpression of AQP-1 in the model and include the effects of increased MMP activity, we increased cell speed, filopodia were stabilized and decreased in number from three to two, and the leaders generated tunnels for the followers. In this scenario, our model predicted that the stream is unlikely to break up and invasion is very robust (compare Model 2 with Model 11) (Fig. 7D, G). Overall, these simulations support the experimental observations that AQP-1 influences cell migration and invasion in multiple ways (Fig. 7H; Supplemental Movie 5).

## DISCUSSION

Our discovery of high AQP-1 expression within lead cells of the chick cranial neural crest cell migratory stream (1, 2) led us to examine the function of this water channel protein. We initially confirmed that *AQP-1* mRNA and protein are both higher in lead cranial neural crest cells in vivo, and we used state-of-the-art imaging to show that AQP-1 expression is localized to neural crest cell membranes including cell filopodia. We tested the hypothesis that AQP-1 regulates cell migration to promote neural crest cell invasion using gain- and loss-of-function of AQP-1 and quantification of cell dynamics obtained from time-lapse imaging sessions. We elucidated four important aspects of AQP-1 function that support its critical in vivo role in neural crest cell migration.

First, we discovered that modulation of AQP-1 expression altered neural crest cell motility and invasive ability. An increase in AQP-1 expression in premigratory cranial neural crest cells resulted in higher cell speeds in vitro and enhanced invasion in vivo (Figs. 2, 3). In contrast, the knockdown of AQP-1 function by MO transfection or microinjection of the chemical blocker AZA into the neural crest cell migratory pathway resulted in slower cell speeds in vitro and reduced invasion in vivo (Figs. 2, 3). These results are consistent with previous data that implicated the role of AQP-1 in cell migration and invasion across a wide variety of adult mouse and human cancer cell types (Saadoun et al., 2005; Chen et al., 2012; Chen et al., 2015; Wei and Dong, 2015; Xiong et al., 2017; Cao et al., 2006; Hu and Verkman, 2006; Klebe et al., 2015).

Second, we found that AQP-1 promoted neural crest cell invasive ability by stabilizing filopodia, supporting a function for AQP-1 in enabling cells to bulldoze through the embryonic microenvironment. Initially, we hypothesized that AQP-1 functions to promote neural crest cell invasion by: (1) rapidly changing the cell volume and shape to permit the cell to infiltrate between gaps in loosely connected mesoderm and dense ECM through which cells travel, or; (2) stabilizing cell filopodia to allow break down and displace the surrounding tissue. In support of (2), we found a significant reduction in the number and length of neural crest cell filopodia after AQP-1 knockdown (Fig. 4). Further, neural crest cell filopodia retracted faster and survived for shorter duration in cells with reduced AQP-1 function (Fig. 4).

We did not observe overall cell volume changes in migrating neural crest cells in vivo by dynamic analysis of three-dimensional cell volumes measured from time-lapse imaging sessions (data not shown). Given that changes in cell size that are induced by alterations in external osmolarity have been linked to AQP-1 function in human vascular smooth muscle cells in culture (Shanahan et al., 1999), we cannot completely rule cell volume changes out since changes in the dynamic filopodia volume of migrating neural crest cells may require higher resolution microscopy to detect. Further, a similar volume of water may flow into or out of control and AQP-1 perturbed neural crest cells as perturbations are not localized to distinct regions of individual cells.

Third, we learned that AQP-1 directly influenced neural crest cell adhesion and ECM degradation, suggesting a mechanistic basis for neural crest cell “bulldozing” through the microenvironment. In support of this, we showed that AQP-1 and phosphorylated-FAK were colocalized in migrating neural crest cells, and upon AQP-1 overexpression, less pFAK and integrin B1 puncta were present on cell surfaces (Fig. 5). This implicated AQP-1 in the integrin-mediated focal adhesion signaling pathways previously shown to be important for neural crest cell migration (Desban and Duband, 1997; Desban et al., 2006; Parsons, 2003). Furthermore, it has been previously shown that Aquaporin-1 regulates FAK expression in bone marrow mesenchymal stem cells (Meng et al., 2014). FAK binds directly to integrin B1, and Aquaporin-2 has been shown to internalize integrin B1 (Chen et al., 2012; Lechertier and Hodivala-Dilke, 2012). Therefore, in migrating neural crest cells, less integrin B1 and pFAK on cell surfaces when AQP-1 is overexpressed may indicate increased integrin turnover and less cell adhesion (Fig. 5). AQP-1 perturbation affected the expression and activity of MMPs in migrating cranial neural crest cells when measured by RNAseq profiling, in-gel zymography and degradation assay (Fig. 5). Whether pFAK colocalization with AQP-1 in migrating neural crest cells also plays a role to initiate downstream intracellular signals, including Eph-ephrins, is unclear (Carter et al., 2002; Miao et al., 2000) and will be the focus of future studies. Together, these data clearly demonstrated that AQP-1 promotes neural crest cell invasion by influencing FAK activity, integrin turnover and ECM degradation.

Fourth, we showed that AQP-1 is involved in directed neural crest cell migration, suggesting that AQP-1 is downstream of a cell’s ability to readout guidance factors in the local microenvironment. AQP-1 and EphB receptors were colocalized in the same migrating neural crest cells at both the RNA and protein levels (Fig. 6). Neural crest cell directionality in response to endogenous guidance cues was reduced when AQP-1 was overexpressed both in vitro and in vivo, as the majority of the AQP-1 water channels were no longer downstream of guidance signals (Figs. 3, 6). Further, AQP-1 overexpression led to observed neural crest cell filopodia pointing in random directions (Supplemental Fig. 2). It was not clear whether AQP-1 is enriched in a directional manner, since although there is higher AQP-1 protein expression in lead migrating neural crest cells in vivo (Fig. 1H), our super resolution microscopy that revealed AQP-1 protein expression in neural crest cell filopodia was performed in vitro in the absence of endogenous guidance cues (Fig. 1E). Future experiments that dissect the AQP-1 and EphB signaling relationship and explore other guidance receptors we previously identified by profiling migrating cranial neural crest cells (McLennan et al., 2015) will help to shed light on this.

Lastly, our computational model simulations identified cell speed, filopodia stabilization and ECM degradation as key parameters for controlling neural crest cell invasion in support of AQP-1 function (Fig. 7). When we increased/decreased cell speed to simulate AQP-1 gain-of-/loss-of-function, respectively, we observed the intuitive changes in the maximum distance traveled by cells (Fig. 7; Models 2,3; Supplemental Movie 5). However, unexpected breakdown of the cell migratory stream occurs in silico when the cell speed is increased; a pattern that is not observed in vivo (Fig. 3). Examination of cell speed versus filopodial dynamics revealed that stabilization of filopodia rather than filopodia number or polarity increased the furthest distance traveled by cells (regardless of speed) and reduced the likelihood of stream separation (Fig. 7). Thus, our computational model simulations support the role of AQP-1 in promoting neural crest cell invasion through ECM degradation, stabilization of filopodia and regulation of cell speed.

In summary, our findings implicated four processes of AQP-1 function in neural crest cell migration: modulated cell speed, stabilized filopodia, cell adhesion and ECM degradation, and co-localization with cell guidance receptors. Whether these functions are dependent or independent of AQP-1 functioning as a water channel is currently unknown, however given the vast recent literature on AQP-1 it is likely to be related to its water functions. By stabilizing filopodia, increasing MMP-mediated ECM degradation and controlling adhesions in response to guidance receptors, lead neural crest cells “bulldoze” through the embryonic microenvironment in a directed manner. Later emerging neural crest cells are able to follow and collective migration is maintained. Computational modeling identified cell speed, filopodia stabilization and ECM degradation as key parameters to promote neural crest cell migration. The combination of computational modeling with an in vivo dynamic imaging platform with single cell resolution provides a powerful tool to examine both the upstream regulation of AQP-1 activity and downstream signaling events in directed cell migration. With the established high correlation of AQP-1 expression and cancer cell aggressiveness (Tomita et al., 2017; De leso and Yool, 2018), our data of in vivo AQP-1 function during embryonic neural crest cell migration and mechanistic basis of cell bulldozing offer further details on and illustrate the importance of the function of aquaporins in development and cancer.

## MATERIALS AND METHODS

### Embryos

White Leghorn, fertilized chicken eggs (Centurion Poultry, Inc.) were incubated in a humidified incubator at 38°C to the desired developmental stage (Hamburger and Hamilton, 1951).

### RNAscope and immunohistochemistry (IHC)

RNAscope on whole chick heads was performed as previously described (Morrison et al., 2017b). Briefly, HH13 chick embryos were harvested and fixed in 4% paraformaldehyde for 2 hours. Following a dehydration gradient in methanol, embryos were stored for 4 days at −20°C before RNAscope protocol began. After rehydration, embryos were digested in a 0.1x dilution of protease solution in PBS-Tween for 6 minutes at 22°C with gentle agitation. AQP-1, EphB1 and EphB3 RNAscope probes were designed by Advanced Cell Diagnostics (Newark, CA) against GenBank accession numbers NM_001039453.1, NM_205035.1, XM_422762.4 respectively. AQP-1 was labeled with Cy3, EphB1 with Atto488 and EphB3 with Cy5 dyes. As a control, embryos were processed as described but no probes were used (Supplemental Fig. 1A). After RNAscope, IHC was performed in a 4% molecular biology grade BSA blocking buffer using HNK-1 primary antibody (TIB-200 hybridoma cell line, ATCC Cell Lines) in 1:25 concentration overnight incubation at 4°C. Secondary antibody Alexa Fluor 594 Goat anti-Mouse IgM (A21044, Thermofisher Scientific, Waltham, MA, USA) was used in 1:500 concentration overnight at 4°C. The embryos were then cleared with FRUIT clearing buffer (Hou et al., 2015) at 60% followed by 80% overnight at 4°C.

IHC on cryosections or whole mount chick heads was performed with either Phospho-FAK (Tyr861) (44-626G, ThermoFisher Scientific) at 1:200 concentration, AQP-1 Millipore AB3272 at 1:50 concentration or EphB1/3 (sc-926, discontinued product, Santa Cruz Biotechnology, Dallas, TX, USA). Generally, tissue was fixed in 4% PFA overnight at 4°. After washing in PBS, tissue was blocked with 4% BSA, 0.1% Tween block for 2 hours at room temperature followed by overnight incubation with primary antibodies at 4°. After washing in block, secondary antibodies were applied in 1:500 concentration overnight at 4°. After a final wash with PBS, whole heads were mounted for imaging and cryosections were coverslipped with VECTASHIELD HardSet Antifade Mounting Medium with DAPI (H-1500, Vector Laboratories, Burlingame, CA). The z stacks of the collected images were loaded into Imaris 9.20. The images were segmented using the surface function with HNK-1 setting (settings: grain size at least 0.439 um, largest sphere 1.65 um, threshold automatically detected at upper end, lower end 110, estimated diameter 6-7um, voxels above 1000). Then, cells were selected at the back and front (respectively) of the r4 stream. The mean intensity of the HNK-1 was recorded. The HNK-1 intensity value was normalized from a spot detected outside the cells on the same slide to account for background. The cells intensity was averaged together and plotted on a box plot via Origin for the front and the back of the stream. Secondary antibodies: pFAK – Goat anti rabbit IgG (H+L) (CM405F, Biotium, Fremont, CA, USA), AQP-1 – F(ab’)2-Goat anti-Rabbit IgG (H+L) Cross-Adsorbed Secondary Antibody, Alexa Fluor 546 (A-11071, ThermoFisher Scientific), EphB1/3 – Alexa Fluor 488 Goat anti-Rabbit 488 (A-11008, ThermoFisher Scientific), HNK-1 – Alexa Fluor 647 Goat anti-Mouse IgM (A-21238, discontinued product, ThermoFisher Scientific). Pearson’s coefficients for colocalized fluorescence were calculated in Imaris 9.0.0 using the Coloc module on 3D data sets. Images of pFAK, HNK-1 and AQP-1 IHC from different regions of the r4 migratory neural crest stream were created using a 40x 1.2NA objective with Airyscan imaging on a LSM 800 (Zeiss, Jena, Germany). Post Airyscan processing, a mask was created with the HNK-1 channel to isolate neural crest and thresholds for pFAK and AQP-1 channels were created to eliminate background. The Pearson’s coefficient for n=5 embryos was recorded and averaged.

### In vitro assays

In vitro neural tube cultures were performed as previously described (McLennan et al., 2010), using Ham’s F-12 Nutrient Mix media (11765054, ThermoFisher Scientific). For the initial experiments to examine cell speed, DMSO was added to the media for controls for the AZA (A6011, Sigma-Aldrich, St Louis, MO) experiments because AZA was solubilized in DMSO. To transfect some cells in the neural tube cultures with AQP-1 FL, dorsal neural tubes were electroporated in ovo as previously described (McLennan and Kulesa, 2007) and allowed to recover for 2-4 hours before being removed and plated in vitro. Alternatively, neural tubes were removed from embryos and then constructs or morpholinos were electroporated into the tissue using petri dish platinum electrode for tissues chamber (45-0505, BTX, Holliston, MA, USA), 5 pulses, 60V. Neural tubes were allowed to recover in chamber for 10 minutes and were then plated. AQP-1 FL construct was designed and built in the lab, using pMES as a backbone. Prior to neural crest experiments, AQP-1 FL was tested in a chick cell line and shown to overexpress AQP-1 (Supplemental Fig. 1B, C). For the directionality towards the BA2 tissue experiments, neural tubes were soaked in Hoechst, 1:200 dilution for 5 minutes prior to plating and BA2 tissue was soaked in DiI (V22889, Thermofisher Scientific), 1:30 dilution for 30 minutes. Time-lapses of cultures were taken on a LSM 710, LSM 780 or LSM 800 (Zeiss), 2.5 minute intervals, single z-planes. For IHC on in vitro cultures, the same antibodies and protocols were followed as for cryosections and whole mount embryos, however the incubation times were shorter. In vitro cultures were fixed with 4% paraformaldehyde for 20 minutes. Cultures were incubated with primary and secondary antibodies for 1 hour each. Images were taken using confocal microscopy and Airyscan detection with a 40x 1.2NA water immersion objective. Due to the lengthy timing of collecting Airyscan z stacks and to reduce possible photobleaching, this imaging was not performed for the entire cell but focused on the cell membranes against the fibronectin-coated dish surface.

The z stacks of the collected images were loaded into Imaris 9.20. The cells were segmented using the surface function to manually outline the shape of the cell. Then, spots were detected on each the integrin and pFAK channels. The integrin channel spots included a diameter of .6um spot with background subtraction and a quality filter was picked using a statistical upper and lower threshold break in the data. The pFAK spots were detected using .4um spots and background subtraction with the quality filter was picked using a statistical upper and lower threshold break in the data. The number of spots for each cell was recorded for pFAK and integrin channels then an average was calculated for each data set. The data was then plotted on a box plot via Origin. For the neural tube cultures with DMSO, AZA or AQP-1 FL, cells were manually tracked using “Spot” detection in Imaris for ≥ 6 hours. The spots were set to ≥ 9 um in size. The mean speed and displacement of each track was calculated using Imaris (Bitplane). Straightness equals displacement of track divided by length of track. The box plots in each figure were generated by using the values from each dataset indicated. X indicates outliers, and the box plots and whiskers indicate the quartiles and range, respectively, of each dataset. P-values were calculated using a standard Student’s *t* test or paired *t* test. Data distribution was assumed to be normal, but this was not formally tested.

### In ovo injections, electroporations and analysis

AZA (500 uM) was injected into multiple sites of the mesoderm on one side adjacent to the hindbrain region of HH8-9 embryos. For control injections, the same injections were performed using DMSO. After 24 hours reincubation, embryos were fixed in 4% paraformaldehyde for 2 hours at room temperature and then immunohistochemistry was performed for HNK-1 as described above. Each cranial region of injected embryos was then cut down the midline and each half mounted in a glass slide as previously described (Teddy and Kulesa, 2004). Both halves of the cranial region of each embryo were imaged on a LSM 800 (Zeiss) so that injected and control halves of each embryo could be compared. In ovo electroporations were performed as previously described (McLennan and Kulesa, 2007). Fluorescein-tagged AQP-1 and control morpholinos were designed by, and obtained from, Gene Tools, LLC (Philomath, OR, USA). H2B mCherry (2.5ug/ul) and morpholinos (0.5mM) were injected and electroporated at HH8-9 and reincubated for 20 hours before being fixed, processed for HNK-1 and imaged as described above. An empty EGFP vector, pMES, was used as control to AQP-1 FL, as AQP-1 FL was inserted into the pMES vector. pMES and AQP-1 FL were either injected and electroporated with DiI before being reincubated for 12 hours or injected and electroporated without DiI before being reincubated for 16 hours. Time-lapses were performed as previously described (McKinney et al., 2013). We calculated the percentage of area covered using the “Surfaces” function of Imaris (Bitplane) to create a surface mask by manually drawing the outline of the whole branchial arch. Next, we calculated the area of the HNK-1 fluorescence signal using the masked arch surface. We set a consistent intensity threshold to the same value for each dataset, a surface grain size of 1 um was set, the diameter of the largest sphere was set to 1 µm, and then the automatic “Surfaces” function was applied. We calculated the percentage of the front of arch the HNK-1 signal covered by comparing the two values in the front 50% of the stream. For electroporated embryos, cells were automatically detected using the “Spot” function in Imaris. The spots were set to ≥ 9 µm in size. The spots were counted in the front 50% of the stream. The percentage of total spots versus cells in the whole stream was calculated. The percentage of distance the cells migrated was calculated by measuring the total distance of the migratory route and measuring what distance the transfected cells migrated. This was calculated on 3D z-stacks using the measurement tool in Imaris. The box plots in each figure were generated by using the values from each dataset indicated. X indicates outliers, and the box plots and whiskers indicate the quartiles and range, respectively, of each dataset. P-values were calculated using a standard Student’s *t* test or paired *t* test. Data distribution was assumed to be normal, but this was not formally tested.

### Spinning Disk Imaging and Analysis

HH13 chick embryos were mounted dorsal side down on glass bottomed dishes with a grease-sealed Teflon membrane over the top to preserve humidity (Rupp and Kulesa, 2007). Samples were placed inside a heated chamber around a spinning disk confocal microscope (PerkinElmer, Ultraview), allowed to acclimate to the chamber, and 2 channel imaging proceeded until any sign of phototoxicity was observed. Cells in the front 10% of the migratory r4 neural crest stream were chosen for imaging. Images were collected with a 40x 1.2NA water immersion objective in 1 um Z-steps for up to 15 um total depth in either 20 or 30 second increments. Time-lapse data were analyzed in a combination of Imaris (Bitplane AG) and ImageJ. If cells were overlapped in Z, Imaris 3D imaging was used to create a mask of the cell of interest which was then projected onto 2D. A projected image of the cell was imported into ImageJ and after smoothing and background subtraction, an 8-bit binary mask was created using the Gap-mTurquoise2 fluorescence. Minor adjustments to the mask were made by hand for extremely thin filopodia or touching cells. The time-lapse masked image was imported into a modified version of the CellGeo software (Tsygankov et al., 2014) that did not use the java enabled GUI but allowed for parameter adjustment using the same algorithms. BisectoGraph and FiloTrak modules were used and parameters of 20 smooth, 1 CrR, 1 CutOff, 9 Critical length, 5.5 critical width were used. Filopodia were tracked manually. The remainder of analysis was written in MATLAB (Mathworks Inc) to collect filopodia data from multiple cells under each condition.

### Cytometry, RNA-seq and Analysis

Premigratory neural crest were electroporated with either AQP-1 FL or pMES and eggs re-incubated for 24 hours. The neural crest stream adjacent in rhombomere 4 was isolated from healthy ∼HH15 embryos. Five pMES transfected embryos were pooled for each of the 3 biological replicates (n=15 total). Four AQP-1 FL embryos were pooled for each of the 3 biological replicates (n=12 total). Tissue was dissociated as previously described (McLennan et al., 2015a). Cells were isolated by FACS, which included forward scatter, side scatter, pulse width, live/dead stain (7AAD) and YFP gates as previously described (McLennan et al., 2015a; Morrison et al., 2017a). Cells were sorted directly into 7ul of Clontech lysis solution containing 0.05% RNAse inhibitor. Following lysis for 5 minutes at room temperature, lysates were immediately frozen on dry ice and stored at −80C. Bulk RNA-seq lysates were thawed on ice. cDNA synthesis and library preparation were performed with SMART-seq v4 Ultra Low Input RNA-seq (634891, Takara, Kusatsu, Shiga, Japan) and Nextera XT DNA sample prep and indexing library preparation kits as recommended by the manufacturer (FC-131-2001, FC-131-2004, and FC-131-1096, Illumina, San Diego, CA). Resulting short fragment libraries were checked for quality and quantity using a Bioanalyzer (Agilent) and Qubit Fluorometer (Life Technologies). Libraries were pooled, re-quantified and sequenced as 50bp single reads on 2 lanes of the Illumina HiSeq 2500 in High Output mode using HiSeq Control Software v2.2.58 (illumine). Following sequencing, Illumina Primary Analysis version RTA v1.18.64 and bcl2fastq2 v2.18 were run to demultiplex reads for all libraries and generate FASTQ files. More than 3 million total alignments were produced per sample. Single end 51-base reads were aligned to the chicken genome galGal4 from UCSC with annotations from Ensembl 84 using STAR (2.5.2b) with options –alignEndsType EndToEnd and sjdbScore 2. Downstream analysis was done in R (3.4.1). Differential expression analysis was performed using the edgeR package (3.18.1). Genes were denoted as differentially expressed if they had a p-value less than .05 and a fold change greater than 1.5-fold (absolute value). Gene ontology enrichment was done using a hypergeometric test on lists of differentially expressed genes. The data discussed in this publication have been deposited in NCBI’s Gene Expression Omnibus and are accessible through GEO Series accession number GSE121131 (https://www.ncbi.nlm.nih.gov/geo/query/acc.cgi?acc=GSE121131).

### In gel zymography

To achieve high levels of MMP secretion from neural crest cells grown in culture, neural tubes were isolated, plated onto the bottom of 4 well dishes (176740, ThermoFisher Scientific) with no coating, covered in 250 ul of Ham’s F-12 media. N=39 neural tubes exposed to 100 uM AZA, n=39 neural tubes exposed to DMSO, 0.05 ul in 1 mL, as control for AZA, n=32 neural tubes exposed to 50 umol/L GM6001 (Anderson et al., 2006), n=32 neural tubes exposed to DMSO, 1 ul in 1 mL, as control for GM6001. After 24 hours of incubation, the media was harvested. Protein in media was concentrated using 3K Pierce concentrator (88512, ThermoFisher Scientific), and then used in gel zymography following suppliers protocol (ZY00100BOX, LC2670, LC2671, LC2675, LC2676, Thermofisher Scientific). The substrate was gelatin. Importantly, an image of the gel prior to colloidal blue staining (LC6025, ThermoFisher Scientific) was taken so that the protein ladder could be clearly seen for determining protein size after staining. Quantification of the bands on the gel image were performed using ImageJ and a protocol from Protocol Place (http://protocol-place.com/assays/gelatin-zymography/).

### Degradation assay

Degradation assay was performed using QCM Gelatin Invadopodia Assay (ECM671, Millipore, Burlington, MA, USA). Following the manufacturers protocol, Cy3-labeled gelatin coated wells in 8-well chambered slides (80821, Ibidi, Martinsried, Germany). The protocol was slightly modified by heating all the gelatin to 60 degrees Celsius so that there was no issue in room temperature and warm gelatin not mixing fully. To optimize cell adhesion, fibronectin was then added over the gelatin as previously described (Garmon et al., 2018). 15 neural tubes (transfected with either pMES or AQP-1 FL) were plated into each well with 500ul of media and incubated for 40 hours. For controls, some wells received no neural tubes, but were still incubated for 40 hours with 500ul of media. Cultures were fixed with 4% paraformaldehyde for 20 minutes and then imaged. MMP2 and MMP9 are secreted proteases and possible gelatin degradation will occur throughout each well, not just where neural crest cells are migrating. Therefore, Cy3-labeled gelatin was imaged at 5 randomly chosen areas of each well using a 40x 1.2NA objective, n=3 wells per condition. The z-stack images were loaded into Imaris 9.20. A surface rendering was created on the whole Cy3 gelatin channel throughout the whole image with the automatic settings used to detect the surface using voxel totals above 10. The average intensity across the whole image for the Cy3 channel was recorded. An average was taken from each condition and a box plot was made in Origin.

## ACKNOWLEDGEMENTS

PMK would like to acknowledge kind and generous funding from the Stowers Institute for Medical Research. We also thank members of the Microscopy, Histology and Molecular Biology core facilities and our Scientific Illustrator, Mark Miller, at the Stowers Institute for Medical Research.

## AUTHOR CONTRIBUTIONS

R McLennan performed in vitro and in vivo functional perturbations of AQP-1, isolation of electroporated cells for FACS and RNA-seq, in vivo AQP-1 and Eph IHC, in vitro pFAK and integrin B1 IHC after AQP-1 FL or pMES transfection, degradation assays and corresponding imaging. MC McKinney performed RNAscope experiments, filopodia dynamics, pFAK IHC on wildtype neural crest cells and analysis on these experiments. DA Ridenour and CA Manthe built and tested the AQP-1 full length construct. JM Teddy performed the image analysis and measurements. JA Morrison and R McLennan performed cell dissociation and analysis of RNA-seq data. JC Kasemeier and R McLennan performed the in gel zymography experiments. JM Teddy, DA Ridenour and CA Manthe performed the initial time-lapses. R Giniunaite performed the modeling simulations under the supervision and guidance of M Robinson, RE Baker and PK Maini. All authors discussed and interpreted the results. R McLennan and PM Kulesa wrote the manuscript and generated the figures with critical comments and suggestions from JA Morrison, JM Teddy and MC McKinney. PM Kulesa supervised the overall project. All authors read and approved the final manuscript.

---

**Supplemental Figure 1.** (A) No probes control for RNAscope experiment. (B) Quantification of AQP-1 RNA expression in chick embryonic fibroblasts (CEF), chick liver hepatocellular carcinoma (LMH) and LMH transfected with AQP-1 FL. (C) Quantification of AQP-1 RNA expression in LMH cells after mock transfection, pMES transfection and AQP-1 FL transfection. (D) Box plot of the Speed (microns/hr) of neural crest cells, transfected with pMES control vector (black, n= 27 cells from n=9 neural tube explants), non-transfected but in the same cultures as pMES (blue, n=26 cells) and transfected with AQP-1 FL in different cultures but prepared and imaged the same days as controls (red, n=33 cells from n=10 neural tube explants). (E) Box plot of the distance migrated by neural crest cells after mesodermal injections of AZA (red) and the distance migrated by neural crest cells on the control sides of the same embryos (black), n=6 embryos. (F) Box plot of the percentage of neural crest cells that migrate into the branchial arches, n=8 embryos per treatment. (G) The entire zymogram that is quantified in Figure 5. An image of the ladder (left) was taken prior to development so that it could be clearly seen. Proteins in these assays do not run exactly the same as the markers as enzymes in the samples are not reduced while the markers are (Woessner, 1995). The band size is approximately 62 kD, corresponding to MMP2 (Anderson, 2010).

**Supplemental Figure 2: Filopodia direction and length with respect to neural crest migratory direction**. (A) Distribution of angles of filopodia for control (black), AQP-1 FL overexpression vector (green) and the AQP-1 MO (red) labeled cells. Radial magnitude equals number of filopodia in that direction. (B) Length and direction of the filopodia for cells labeled with control vector, AQP-1 FL overexpression vector or AQP-1 morpholino. Radial magnitude equals length of the filopodia in microns and angle is with respect to migratory direction. Contrasting colored filopodia are in a 30° window around the direction of migration. (C) Example neural crest filopodia with direction of migration indicated. Direction of migration is always set to 0 degrees.

**Supplemental Figure 3:** Schematic of a rectangular domain we used to model the domain on which the neural crest cells migrate (black circles). They enter the domain from the neural tube (x = R). BA2 denotes branchial arch 2.

**Supplemental Table 1:** Genes that are differentially expressed between pMES (control) and AQP-1 FL. (excel document, not attached)

**Supplemental Table 2: Model parameters used in the simulations provided in the results section.**

**Supplemental Movie 1: Cranial neural crest cell migratory behaviors are altered in vitro after AQP-1 manipulation.** (top) Control migrating neural crest cells exposed to DMSO. (middle) AQP-1 was inhibited by adding Acetazolamide (AZA) to the media. AZA was solubilized in DMSO. (bottom) AQP-1 was overexpressed by transfection of AQP-1 full length construct (green cells). Cranial neural tube explants are shown on the left-hand side of each panel. Time intervals between images ranged from 2.5 to 4 minutes and frame speed was adjusted so that each time-lapse was 8 hours in duration. Scale bar= 30 um.

**Supplemental Movie 2: Cranial neural crest cell migratory behaviors are altered in vivo after AQP-1 manipulation.** (Left) Premigratory neural crest cells were transfected with Gap43-YFP/H2B mCherry (control). (Right) AQP-1 full length/H2B mCherry. Z-stacks were collected every 5 minutes and approximately 6 hours is shown. Scale bar= 20 um.

**Supplemental Movie 3: Fast confocal imaging reveals changes in neural crest cell filopodial dynamics after AQP-1 manipulation.** Projected images from spinning disk time-lapse microscopy of migrating <u>lead</u> neural crest cells in whole embryo culture electroporated with either (top) pMES control, (middle) AQP-1 FL or (bottom) AQP-1 Morpholino (MO). Each movie sequence shows the cell membrane label (Gap43-mTurquoise2) to highlight the cell protrusion dynamics. Images were collected in 30 second intervals and shown here for approximately 11 min.

**Supplemental Movie 4: Cranial neural crest cell directionality is altered after AQP-1 manipulation.** Premigratory neural crest were transfected with pMES (control) or AQP-1 FL and neural tubes were plated in the presence of branchial arch 2 (ba2) tissue as a source of known endogenous chemoattraction. Images were collected every 5 minutes and a total of approximately 18 hours of elapsed time is shown. Scale bar= 50 um.

**Supplemental Movie 5: Computer model simulations of cranial neural crest cell migration with AQP-1 manipulation.** (top) Control migration is modeled by normal cell speed and unstable filopodia. (middle) AQP-1 loss-of-function is modeled by reduced cell speed and reduced number of cell filopodia. (bottom) AQP-1 gain-of-function is modeled by increased cell speed, stable cell filopodia and tunneling. Each simulation is run on a 2D migratory domain.

## REFERENCES

Agre, P, Sasaki, S, Chrispeels, MJ. Aquaporins: a family of water channel proteins. Am J Physiol. 1993 265(3 Pt 2):F461.

Ameli, PA, Madan, M, Chigurupati, S, Yu, A, Chan, SL, Pattisapu, JV. Effect of acetazolamide on aquaporin-1 and fluid flow in cultured choroid plexus. Acta Neurochir Suppl. 2012 113:59–64.

Anderson, RB, Turner, KN, Nikonenko, AG, Hemperly, J, Schachner, M, Young, HM. The cell adhesion molecule l1 is required for chain migration of neural crest cells in the developing mouse gut. Gastroenterology. 2006 130(4):1221–32.

Bin, K, Shi-Peng, Z. Acetazolamide inhibits aquaporin-1 expression and colon cancer xenograft tumor growth. Hepatogastroenterology. 2011 58(110-111):1502–6.

Cai, L, Chen, WN, Li, R, Hu, CM, Lei, C, Li, CM. Therapeutic effect of acetazolamide, an aquaporin 1 inhibitor, on adjuvant-induced arthritis in rats by inhibiting NF-κB signal pathway. Immunopharmacol Immunotoxicol. 2018 40(2):117–125.

Cao, C, Sun, Y, Healey, S, Bi, Z, Hu, G, Wan, S, Kouttab, N, Chu, W, Wan, Y. EGFR-mediated expression of aquaporin-3 is involved in human skin fibroblast migration. Biochem J. 2006 400(2):225–34.

Carter, N, Nakamoto, T, Hirai, H, Hunter, T. EphrinA1-induced cytoskeletal re-organization requires FAK and p130(cas). Nat Cell Biol. 2002 4(8):565–73.

Chen, Y, Rice, W, Gu, Z, Li, J, Huang, J, Brenner, MB, Van Hoek, A, Xiong, J, Gundersen, GG, Norman, JC, Hsu, VW, Fenton, RA, Brown, D, Lu, HA. Aquaporin 2 promotes cell migration and epithelial morphogenesis. J Am Soc Nephrol. 2012 23(9):1506–17.

Chen, J, Wang, Z, Xu, D, Liu, Y, Gao, Y. Aquaporin 3 promotes prostate cancer cell motility and invasion via extracellular signal-regulated kinase 1/2-mediated matrix metalloproteinase-3 secretion. Mol Med Rep. 2015 11(4):2882–8.

Condeelis, J. Life at the leading edge: the formation of cell protrusions. Annu Rev Cell Biol. 1993 9:411–44.

Cowan, CA, Yokoyama, N, Bianchi, LM, Henkemeyer, M, Fritzsch, B. EphB2 guides axons at the midline and is necessary for normal vestibular function. Neuron. 2000 26(2):417–30.

De leso, ML, Yool, AJ. Mechanisms of Aquaporin-Faciliated Cancer Invasion and Metastasis. Front Chem. 2018 6:135.

Desban, N, and Duband, JL. Avian neural crest cell migration on laminin: interaction of the alpha1beta1 integrin with distinct laminin-1 domains mediates different adhesive responses. J Cell Sci. 1997 110(Pt 21):2729–44.

Desban, N, Lissitzky, JC, Rousselle, P, Duband, JL. alpha1beta1-integrin engagement to distinct laminin-1 domains orchestrates spreading, migration and survival of neural crest cells through independent signaling pathways. J Cell Sci. 2006 119(Pt 15):3206–18.

Ding, T, Ma, Y, Li, W, Liu, X, Ying, G, Fu, L, Gu, F. Role of aquaporin-4 in the regulation of migration and invasion of human glioma cells. Int J Oncol. 2011 38(6):1521–31.

Garmon, T, Wittling, M, Nie, S. MMP14 Regulates Cranial Neural Crest Epithelial-to-Mesenchymal Transition and Migration. Dev Dyn. 2018 247(9):1083–1092.

Gustafsson MGL, Shao L, Carlton PM, Wang, CLR, Golubovskaya, IN, Cande, WZ, Agard, DA, Sedat, JW. Three-Dimensional Resolution Doubling in Wide-Field Fluorescence Microscopy by Structured Illumination. Biophys. J. 2008 94:4957–4970.

Hamburger, V, Hamilton, HL. A series of normal stages in the development of the chick embryo. J Morphol. 1951 88(1):49–92.

Hou, B, Zhang, D, Zhao, S, Wei, M, Yang, Z, Wang, S, Wang, J, Zhang, X, Liu, B, Fan, L, Li, Y, Qiu, Z, Zhang, C, Jiang, T. Scalable and DiI-compatible optical clearance of the mammalian brain. Front Neuroanat. 2015 9:19.

Hu, J, Verkman, AS. Increased migration and metastatic potential of tumor cells expressing aquaporin water channels. FASEB J. 2006 20(11):1892–4.

Huber, VJ, Tsujita, M, Yamazaki, M, Sakimura, K, Nakada, T. Identification of arylsulfonamides as Aquaporin 4 inhibitors. Bioorg Med Chem Lett. 2007 17(5):1270–3.

Ishibashi, K, Kondo, S, Hara, S, Morishita, Y. The evolutionary aspects of aquaporin family. Am J Physiol Regul Integr Comp Physiol. 2011 300(3):R566–76.

Jiang, B, Li, Z, Zhang, W, Wang, H, Zhi, X, Feng, J, Chen, Z, Zhu, Y, Yang, L, Xu, H, Xu, Z. miR-874 Inhibits cell proliferation, migration and invasion through targeting aquaporin-3 in gastric cancer. J Gastroenterol. 2014 49(6):1011–25.

Karlsson, T, Bolshakova, A, Magalhães, MA, Loitto, VM, Magnusson, KE. Fluxes of water through aquaporin 9 weaken membrane-cytoskeleton anchorage and promote formation of membrane protrusions. PLoS One. 2013 8(4):e59901.

Klebe, S, Griggs, K, Cheng, Y, Driml, J, Henderson, DW, Reid, G. Blockade of aquaporin 1 inhibits proliferation, motility, and metastatic potential of mesothelioma in vitro but not in an in vivo model. Dis Markers.2015 2015:286719.

Lechertier, T, Hodivala-Dilke, K. Focal adhesion kinase and tumour angiogenesis. J Pathol. 2012 226(2):404–12.

McKinney, MC, Fukatsu, K, Morrison, J, McLennan, R, Bronner, ME, Kulesa, PM. Evidence for dynamic rearrangements but lack of fate or position restrictions in premigratory avian trunk neural crest. Development. 2013 140(4):820–30.

McLennan, R, Bailey, CM, Schumacher, LJ, Teddy, JM, Morrison, JA, Kasemeier-Kulesa, JC, Wolfe, LA, Gogol, MM, Baker, RE, Maini, PK, Kulesa, PM. DAN (NBL1) promotes collective neural crest migration by restraining uncontrolled invasion. J Cell Biol. 2017 216(10):3339–3354.

McLennan, R, Dyson, L, Prather, KW, Morrison, JA, Baker, RE, Maini, PK, Kulesa, PM. Multiscale mechanisms of cell migration during development: theory and experiment. Development. 2012 139(16):2935–44.

McLennan, R, Kulesa, PM. In vivo analysis reveals a critical role for neuropilin-1 in cranial neural crest cell migration in chick. Dev Biol. 2007 301(1):227–39.

McLennan, R, Schumacher, LJ, Morrison, JA, Teddy, JM, Ridenour, DA, Box, AC, Semerad, CL, Li, H, McDowell, W, Kay, D, Maini, PK, Baker, RE, Kulesa, PM. Neural crest migration is driven by a few trailblazer cells with a unique molecular signature narrowly confined to the invasive front. Development. 2015a 142(11):2014–25.

McLennan, R, Schumacher, LJ, Morrison, JA, Teddy, JM, Ridenour, DA, Box, AC, Semerad, CL, Li, H, McDowell, W, Kay, D, Maini, PK, Baker, RE, Kulesa, PM. VEGF signals induce trailblazer cell identity that drives neural crest migration. Dev Biol. 2015b 407(1):12–25.

McLennan, R, Teddy, JM, Kasemeier-Kulesa, JC, Romine, MH, Kulesa, PM. Vascular endothelial growth factor (VEGF) regulates cranial neural crest migration in vivo. Dev Biol. 2010 339(1):114–25.

Meng, F, Rui, Y, Xu, L, Wan, C, Jiang, X, Li, G. Aqp1 enhances migration of bone marrow mesenchymal stem cells through regulation of FAK and β-catenin. Stem Cells Dev. 2014 23(1):66–75.

Miao, H, Burnett, E, Kinch, M, Simon, E, Wang, B. Activation of EphA2 kinase suppresses integrin function and causes focal-adhesion-kinase dephosphorylation. Nat Cell Biol. 2000 2(2):62–9.

Morrison, JA, McKinney, MC, Kulesa, PM. Resolving in vivo gene expression during collective cell migration using an integrated RNAscope, immunohistochemistry and tissue clearing method. Mech Dev. 2017b 148:100–106.

Morrison, JA, McLennan, R, Wolfe, LA, Gogol, MM, Meier, S, McKinney, MC, Teddy, JM, Holmes, L, Semerad, CL, Box, AC, Li, H, Hall, KE, Perera, AG, Kulesa, PM. Single-cell transcriptome analysis of avian neural crest migration reveals signatures of invasion and molecular transitions. Elife. 2017a 6. pii: e28415.

Papadopoulos, MC, Saadoun, S, Verkman, AS. Aquaporins and cell migration. Pflugers Arch. 2008 456(4):693–700.

Parsons, JT. Focal adhesion kinase: the first ten years. J Cell Sci. 2003 116(Pt 8):1409–16.

Rupp, PA, Kulesa, PM. High-Resolution, Intravital 4D Confocal Time-Lapse Imaging in Avian Embryos Using a Teflon Culture Chamber Design. CSH Protoc. 2007 2007:pdb.prot4790.

Saadoun, S, Papadopoulos, MC, Hara-Chikuma, M, Verkman, AS. Impairment of angiogenesis and cell migration by targeted aquaporin-1 gene disruption. Nature. 2005 434(7034):786–92.

Shanahan, CM, Connolly, DL, Tyson, KL, Cary, NR, Osbourn, JK, Agre, P, Weissberg, PL. Aquaporin-1 is expressed by vascular smooth muscle cells and mediates rapid water transport across vascular cell membranes. J Vasc Res. 1999 36(5):353–62.

Stroka, KM, Jiang, H, Chen, SH, Tong, Z, Wirtz, D, Sun, SX, Konstantopoulos, K. Water permeation drives tumor cell migration in confined microenvironments. Cell. 2014 157(3):611–23.

Teddy, JM, Kulesa, PM. In vivo evidence for short- and long-range cell communication in cranial neural crest cells. Development. 2004 131(24):6141–51.

Tomita, Y, Dorward, H, Yool, AJ, Smith, E, Townsend, AR, Price, TJ, Hardingham, JE. Role of Aquaporin 1 signaling in cancer development and progression. Int J Mol Sci 2017 18(2). Pii:E299.

Tsygankov, D, Bilancia, CG, Vitriol, EA, Hahn, KM, Peifer, M, Elston, TC. CellGeo: a computational platform for the analysis of shape changes in cells with complex geometries. J Cell Biol. 2014 204(3):443–60.

Verkman, AS. Aquaporins: translating bench research to human disease. J Exp Biol. 2009 212:1707–15.

Wei, X, Dong, J. Aquaporin 1 promotes the proliferation and migration of lung cancer cell in vitro. Oncol Rep. 2015 34(3):1440–8.

Woessner, JF Jr. Quantification of matrix metalloproteinases in tissue samples. Methods Enzymol. 1995 248:510–28.

Xiong, G, Chen, X, Zhang, Q, Fang, Y, Chen, W, Li, C, Zhang, J. RNA interference influenced the proliferation and invasion of XWLC-05 lung cancer cells through inhibiting aquaporin 3. Biochem Biophys Res Commun. 2017 485(3):627–634.

Xu, H, Xu, Y, Zhang, W, Shen, L, Yang, L, Xu, Z. Aquaporin-3 positively regulates matrix metalloproteinases via PI3K/AKT signal pathway in human gastric carcinoma SGC7901 cells. J Exp Clin Cancer Res. 2011 30:86.

Zhang, J, An, Y, Gao, J, Han, J, Pan, X, Pan, Y, Tie, L, Li, X. Aquaporin-1 translocation and degradation mediates the water transportation mechanism of acetazolamide. PLoS One. 2012 7(9):e45976.

